# Mapping the evolution of computationally designed protein binders

**DOI:** 10.1101/2025.10.04.680454

**Authors:** Miguel A. Alcantar, Alexandra M. Paulk, Shoeib Moradi, Debjani Bhar, Grant L. J. Keller, Tanmoy Sanyal, Hua Bai, Gamze Camdere, Seog Joon Han, Mani Jain, Brandon Jew, Sezen Vatansever Inak, Christopher J. Langmead, Christine E. Tinberg, Irwin Chen, Chang C. Liu

## Abstract

Computational protein design enables the generation of binders that target specific epitopes on proteins. However, current approaches often require substantial screening from which hits require further affinity maturation. Methods for experimentally improving designed proteins and exploring their sequence-affinity landscapes could therefore streamline the development of high-affinity binders and inform future design strategies. Here, we use OrthoRep, a system for continuous hypermutation *in vivo*, to drive the evolution of computationally designed mini protein binders (“minibinders”) that target a mammalian receptor. Despite their small sizes (59–72 amino acids), we successfully affinity matured multiple minibinders through strong selection for improved binding and also sampled new regions of minibinder fitness landscapes through extensive neutral drift. One evolved minibinder variant was used to construct a combinatorially complete sequence-affinity map for its six affinity increasing mutations, which revealed nearly full additivity in their contributions to binding. Another minibinder was subjected to both deep mutational scanning and extensive evolution under weak selection, resulting in an evolutionarily diverged collection of binder sequences that revealed non-additive relationships among mutations. Our results highlight that the affinity of computationally designed binders can be rapidly increased through evolution and provide a scalable approach for the evolutionary exploration and subsequent mapping of sequence-affinity landscapes. We suggest that this work will complement protein binder design both as a reliable experimental optimization process and as a vehicle for generating new training data.

## Introduction

The ability to forward engineer new protein binders, for example to disrupt native protein-protein interactions or to create synthetic protein complexes and networks, holds immense importance in biotechnology and medicine^1–5^. Traditionally, custom protein binders such as antibodies have originated from animal immunization^6,7^ or high-throughput screening and selection technologies that isolate binders from large protein libraries^8,9^. However, these experimental discovery approaches are limited in their ability to direct binding to a predefined epitope and thus often fall short of achieving binders with the desired biological effect^10^. In contrast, computational protein design offers a solution to epitope-specific binder discovery, as exemplified in recent work demonstrating *de novo* generation of mini protein binders (“minibinders”) and antibodies targeting proteins at user-specified “hotspots”^11–13^. However, success rates and potencies of successes can be low^11,14^, especially when designing complex interfaces (*e.g*., antibody loops^15^) in engaging difficult targets (*e.g*., membrane proteins, hydrophilic surfaces, or interfaces underrepresented in the PDB^16,17^). These shortcomings motivate the practical question of whether computational designs can be easily improved through directed evolution^18^ and the more basic question of how complex the sequence-affinity landscapes governing designed proteins really are. The answers to these two questions may shape the future of binder engineering, for instance by canonizing a hybrid design plus evolution strategy or projecting the need for sequence-function datasets large enough to contain the complexities of sequence-target-affinity landscapes for training new models. In this work, we address these two questions by leveraging a continuous hypermutation platform, OrthoRep, to affinity mature computationally designed minibinders and broadly explore their sequence-function landscape using a novel neutral drift approach that generates large collections of multi-mutant binders through extensive evolutionary divergence from a single parental design. We found that minibinders readily evolved higher affinity under strong evolutionary pressure for improved binding, which also allowed us to map the combinatorial fitness landscape of affinity-increasing mutations. This revealed that residues not directly involved with binding play a substantial role in improving affinity, and that affinity-increasing mutations evolved under persistent strong selection contributed additively (*i.e*., with minimal epistatic interactions). We also found that sequence-affinity relationships contained in multi-mutant minibinder collections resulting from a neutral drift evolution approach reveal context dependences that a deep mutational scan could not predict, providing a view into the complexity of sequence-affinity landscapes for what are nominally simple topology minibinder designs. Together, these strategies demonstrate how evolution-based approaches can be used to explore the sequence landscape of computationally designed protein binders, providing a direct route to their improvement and a broader template to generating datasets for studying protein-protein interactions in general.

## Results

### Computational methods successfully design minibinders against IL-7Rα

Our initial goal was to design minibinders that target the extracellular domain of human IL-7Rα. As the foundation for the minibinder design campaign, we adapted an approach developed by Cao et al^11^. Briefly, we started with generic miniprotein backbone scaffolds and used ProteinMPNN^19^ to generate structures with competent geometries that would favorably interact with the IL-7 binding site of IL-7Rα. Given that the hit rate for functional binders from this approach is generally low (<1%; refs^11,14^), we further filtered the designs for high AlphaFold-multimer model^20,21^ pLDDT scores—indicative of high-confidence structural predictions. The best predicted binders were cloned and screened using yeast display. Following initial screening, minibinders that showed positive binding on yeast were expressed and purified for binding characterization with surface plasmon resonance (SPR). We focused our efforts on four minibinders that exhibited diverse predicted structures and sequences relative to one another (Figure S1 and Table S1). These four binders, designated Mb-1 through Mb-4, were able to bind soluble IL-7Rα with dissociation constants (K_D_) ranging from 7.8 nM to greater than 1000 nM (Figure 1B; top). These results confirmed that our design approach successfully generated new minibinders with modest affinities for IL-7Rα.

**Figure 1.**
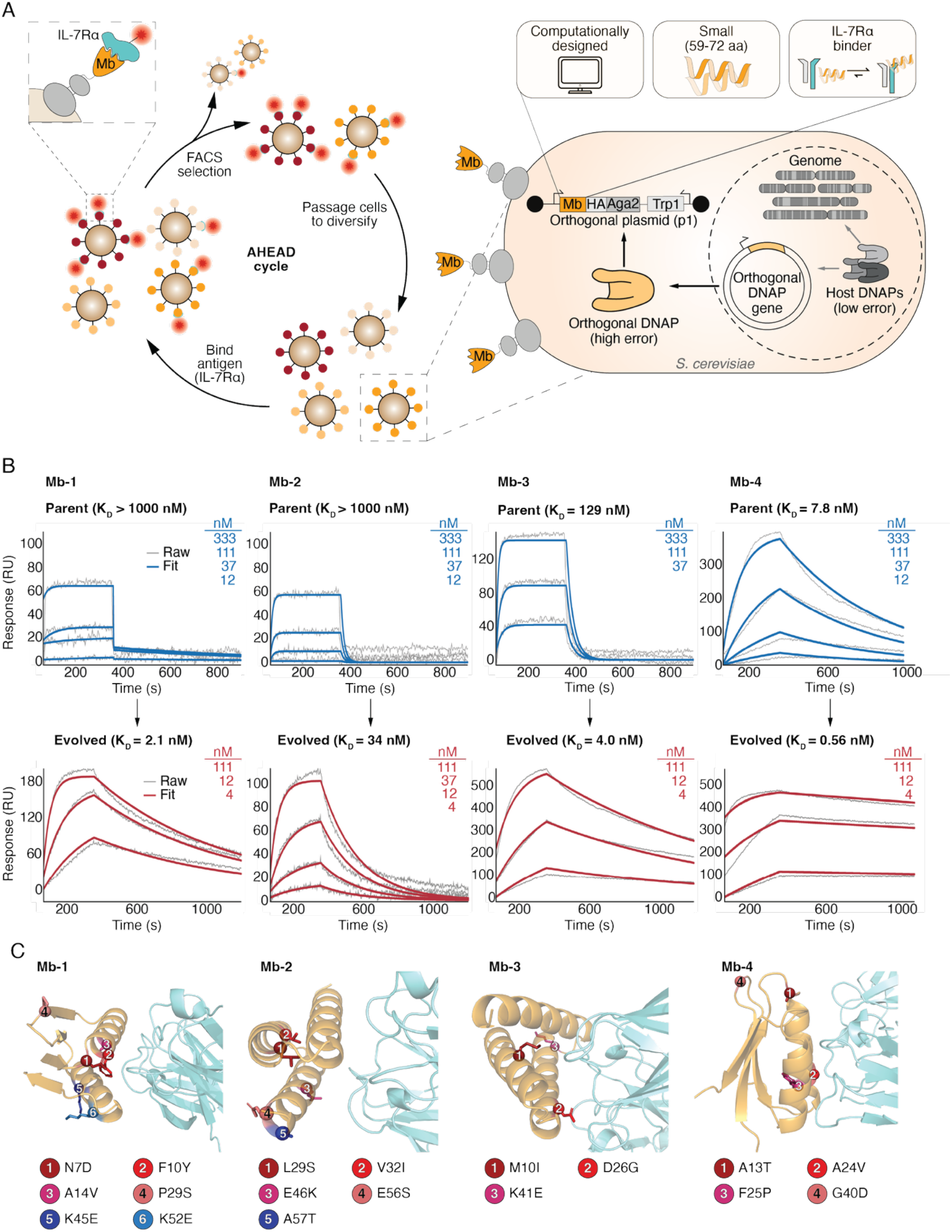
The AHEAD platform enables affinity maturation of minibinder proteins, which are small, computationally-designed proteins that bind the extracellular domain of human interleukin-7 receptor subunit alpha (IL-7Rα). **A**, Schematic overview of the AHEAD platform for rapidly evolving high-affinity minibinders. **B**, Surface plasmon resonance (SPR) traces and associated monovalent affinities for four minibinders before (top) and after (bottom) affinity maturation with AHEAD. The grey lines show raw sensorgram data and the blue (parent) and red (evolved) lines represent curve fits used to extract kinetic parameters. For the evolved variants, the SPR traces are shown for one representative biological replicate and the reported dissociation constants (K_D_) are the mean of two biological replicates (Figure S7). **C**, AlphaFold3 structural predictions for interactions between parent minibinders (orange) and the extracellular domain of IL-7Rα (cyan). The spheres indicate the location of mutations acquired through affinity maturation and the side chains correspond to the parent residue.

### Affinity maturation improves the binding affinities of the initial minibinder designs

To improve the binding affinities of the four computationally designed minibinders, we initiated affinity maturation campaigns using the OrthoRep system^22,23^. OrthoRep is a continuous hypermutation system that uses an error-prone DNA polymerase to mutagenize sequences encoded on a cytosolic plasmid (p1) at an error rate up to ~10^−4^ substitutions per base (s.p.b.), without affecting the genomic mutation rate (~10^−10^ s.p.b.). In one implementation of OrthoRep, protein binders are encoded on p1 and displayed on the yeast cell surface, where they are subjected to selection based on their ability to bind a target protein^24,25^. The iterative cycle of (i) culturing cells to diversify binders using OrthoRep, (ii) displaying and selecting for the best binders, and (iii) passaging to continue mutagenesis of the best binders from the previous selection cycle constitutes the basis of AHEAD (autonomous hypermutation yeast surface display; Figure 1A). Compared to traditional protein binder affinity maturation campaigns that rely on manual cycles of gene diversification (*e.g*., error-prone PCR, targeted combinatorial mutagenesis, or mutation shuffling) and library-scale transformations, AHEAD provides a more streamlined approach for producing high-affinity binders. We have previously used AHEAD to rapidly improve the binding affinities for single domain antibodies^12,24,25^ (nanobodies), and we hypothesized that this workflow could also be used to evolve minibinders, despite their being approximately half the size of nanobodies.

Each of the four minibinders underwent six to nine cycles of AHEAD (Figure S2-S5). During these cycles, we used fluorescence-activated cell sorting (FACS) to select for minibinder-displaying yeast cells exhibiting the strongest binding of IL-7Rα. In most cycles, we successively lowered the concentration of IL-7Rα to increase the selection stringency. At the end of the evolution campaigns, we subcloned a library of sequences from the final AHEAD cycles onto a standard yeast nuclear plasmid and performed two additional rounds of selection to isolate clones expressing the top-performing binders. Dose-response experiments revealed that all four evolved minibinders exhibited enhanced binding relative to their parental counterparts when displayed on the yeast cell surface (Figure S6). This improvement was corroborated using purified proteins, which showed enhanced monovalent K_D_ values ranging from 0.56 to 34 nM (Figure 1B, bottom; Figure S7 and Table S2) and representing 13-to >476-fold improvements in binding affinity relative to the parental variants.

The evolved minibinders each contain three to six mutations relative to their parental sequences (Figure 1C). To examine the structural context of these mutations, we used AlphaFold3 to predict the structures of each minibinder in complex with IL-7Rα. Although AlphaFold-multimer was originally used to filter candidate binders in the design stage, we opted to use the newly available AlphaFold3 to assess structural impacts of mutations in the evolved minibinder variants due to its reported improved performance over AlphaFold-multimer^26^. Additionally, the AlphaFold3 models of the parental minibinders agreed closely with the original design models from AlphaFold2 (Figure S8). We found that while some mutations arose in residues predicted to contact IL-7Rα, many mutations also emerged at positions not expected to contact IL-7Rα based on the structural models (Table S3). For example, in Mb-1, mutations N7D and F10Y were predicted to contact H191 and Y192 on IL-7Rα, respectively. Conversely, K45E and K52E mutations occurred on a face of Mb-1 which appears not to interact with IL-7Rα. Similarly, A14V represents a conversion to a more hydrophobic residue and was buried in the core of Mb-1, making little if any contact with IL-7Rα. The prevalence of affinity-enhancing mutations in residues not directly involved in binding, which has also been observed in other binder scaffolds^18,27^, may contribute to improving intramolecular interactions by fine-tuning the binding interface pose through allostery and/or stabilizing the minibinder structure. Taken together, these results demonstrate that OrthoRep can be used to affinity mature computationally designed minibinders for their cognate target by selecting for beneficial mutations throughout the binder sequence.

### Characterizing a combinatorially-complete library reveals the binding landscape of Mb-1

To more comprehensively investigate how specific mutations observed in evolved minibinder variants contributed to improved binding, we generated a combinatorially complete library of Mb-1. Specifically, we sought to characterize how mutations—both individually and collectively—alter the minibinder’s binding affinity for IL-7Rα. We focused on Mb-1, as its evolved variant exhibited the largest improvement in binding affinity relative to the parental variant (from K_D_ >1000 nM to 2.1 nM) and contained six mutations—the most among the evolved variants. This provided a wide range of binding affinities and mutational combinations to investigate.

We constructed a combinatorially complete library encompassing all 64 possible variants defined by the six mutations observed in the evolved Mb-1 variant: N7D, F10Y, A14V, P29S, K45E, and K52E. To characterize binding, we used a high-throughput method, Tite-seq^28^, to measure the binding affinities of all variants in a pooled format (Figure 2A and Figure S9). Tite-seq is a pooled assay that uses FACS to bin yeast cells displaying protein binders based on binding strength (*e.g*., no, low, medium, and high binding). The distribution of each protein binder variant across these bins is recovered using next-generation sequencing (NGS), and these data are used to reconstruct the relative binding strength of each individual variant. This process is repeated at multiple target concentrations (Figure S9) and the resultant dose-response curves can then be used to obtain an EC_50_ that accurately estimates K_D_ under the assumption that K_D_ is dominated by slow off rates, which is typical for binders.

**Figure 2.**
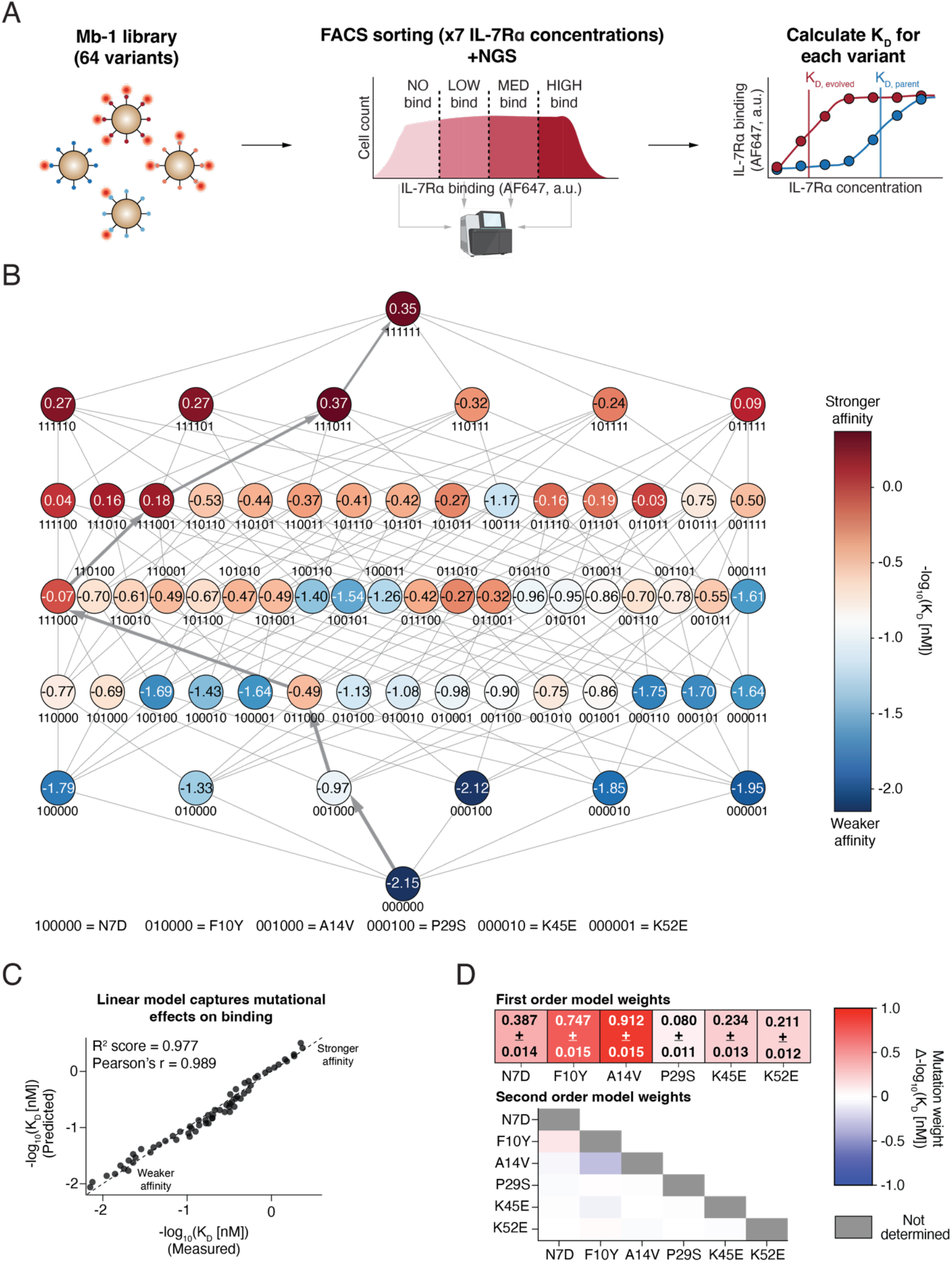
The combinatorial binding landscape between the parental and affinity matured Mb-1 can be described with a linear model. **A**, Schematic overview of the pooled assay, Tite-seq, used to characterize the combinatorial landscape for Mb-1. The Mb-1 variants (64 in total) were sorted into bins based on binding at seven antigen concentrations (0 to 500 nM). The resultant distributions of minibinders across bins were used to reconstruct the activity of individual minibinders and extract their half maximal effective concentrations (EC_50_) to estimate K_D_. **B**, The binding landscape for Mb-1 defined by N7D, F10Y, A14V, P29S, K45E, and K52E. The “greedy” path to the affinity matured Mb-1 is indicated with grey arrows. The values in each circle correspond to the mean −log10(K_D_) for two biological replicates. **C**, A linear model accurately predicts the impact of mutations on binding affinity. The R^2^ score and Pearson’s r are based on the model’s predictions across all 64 variants. The arrows represent the greediest mutational path from the parent to fully evolved variant. **D**, Weights from first-order (linear) and second-order models used to predict the impact of mutations on binding affinity and interactions between mutations, respectively. The mean weights across 100 different train-test splits are shown. For the linear model, values are reported as the mean + one standard deviation.

This pooled assay precisely captured binding affinities across the combinatorial sequence landscape, with strong agreement between two biological replicates (Figure S10; Pearson’s r = 0.911). Using the estimated K_D_ values for all variants, we constructed a combinatorially-complete binding landscape for Mb-1 (Figure 2B). We found that all six mutations individually contributed to improved binding to varying degrees. For example, the P29S mutation, which is located in a β-strand distal to the binding interface, conferred only a modest benefit with regards to improved binding (Δ-log_10_[K_D_] ≈ 0.03). In contrast, the A14V mutation is located on an α-helix proximal to IL-7Rα and provided the largest contribution to improved binding (Δ-log_10_[K_D_] ≈ 1.18).

### A linear biochemical model captures the mutational landscape of Mb-1

To systematically explore how combinations of mutations shape the binding affinity landscape of Mb-1, we used the −log_10_[K_D_] values to construct biochemical models of mutational effects on binding. We began by building a linear model that assumes additive energy effects of individual mutations, estimating the binding affinity of a variant by summing the energy contributions of each mutation. This model captures the mutational landscape without explicitly accounting for possible interactions (*i.e*., epistasis) between mutations. Notably, the linear model captured the binding landscape well, with a representative model achieving a coefficient of determination (R^2^) of 0.977 (Figure 2C). The weights assigned to each mutation—which can be interpreted as the contribution of each mutation toward improving the energy of binding (*i.e*., Δ-log_10_[K_D_])—also strongly agreed with the effects observed in the experimentally measured landscape (Figure 2D, top).

Extending the model to include higher-order interaction terms did not substantially improve predictive performance (Figure S11). However, analysis of the interaction terms from a second-order model, which accounts for pairwise epistasis, revealed a possible synergistic interaction between F10Y and N7D, as well as a potential antagonistic interaction between F10Y and A14V (Figure 2D, bottom). These findings suggest that while some epistatic effects exist, the overall binding landscape defined by mutations in the affinity-matured Mb-1 is largely linear.

### A permissive selection regime enables extensive diversification of a minibinder

The goal of an affinity maturation campaign is to obtain improved binders, which are usually most quickly reached through greedy search where strong positive selection for target binding is applied at each round. This results in frequent selective sweeps, prioritizing enrichment of high-affinity binders at the expense of preserving sequence diversity. Earlier versions of OrthoRep were well-suited for greedy search, as strong selection pressure—enforced by selecting for only the best binders at consecutively lower target concentrations (Figure S2-S5)—could reliably pull evolution along multi-mutation affinity-increasing trajectories even with modest mutation rates underlying diversity generation (~10^−5^ s.p.b.). However, recent advances in the OrthoRep system have increased its mutation rate to ~10^−4^ s.p.b., offering an opportunity to drive extensive protein diversification without relying on the strong pull of stringent positive selection to traverse long mutational paths. As previously shown, this capability is well-suited to broader exploration of sequence-function landscapes^23^.

Leveraging high mutation rate (*i.e*., 10^−4^ s.p.b.) OrthoRep systems, we initiated a protein binder evolution campaign that prioritized functional sequence divergence rather than rapid improvement of affinity—*i.e*., neutral drift. Our approach was based on a modified AHEAD cycle wherein we selected for variants that retained parent-like binding activity (Figure 3A). Successive rounds of this permissive selection strategy were expected to yield a diversified library of binders with comparable binding properties to the parental variant. We focused our diversification campaign on the parental sequence of Mb-4, which had the highest affinity among all parental minibinders. Operating under permissive conditions (*i.e*., high IL-7Rα concentrations relative to K_D_), this choice allowed us to perform many rounds of diversification without requiring prohibitive amounts of purified IL-7Rα. Over the course of 30 AHEAD cycles (Figure S12), we observed a monotonic increase in the average mutation distance from the parent sequence (Figure 3B). After eight cycles, the resultant Mb-4 variants averaged two mutations relative to the parental sequence. By cycle 30, the average sequence contained six mutations which corresponds to an average divergence of approximately 8.3% with respect to the parent sequence.

**Figure 3.**
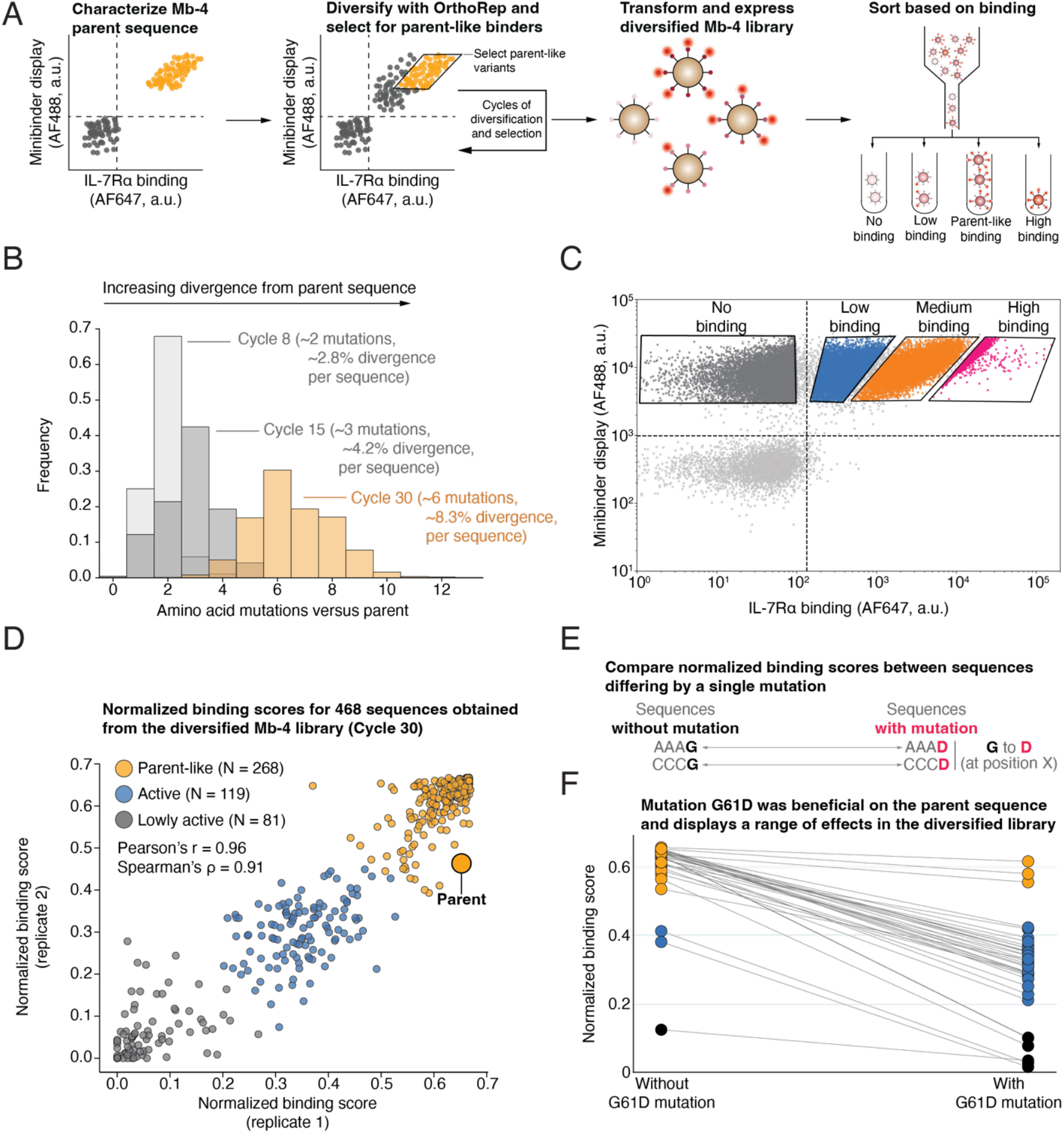
A modified AHEAD cycle can be used to evolve Mb-4 through neutral drift and generate diverse functional minibinder libraries from a parental sequence. **A**, Schematic overview of the modified AHEAD cycle and downstream variant characterization. A parent population is defined during the first cycle and cells that fall within this parent population window are selected during subsequent cycles. This workflow is intended to diversify the minibinder while biasing for functional variants. After several cycles, the diversified library is characterized by FACS sorting based on antigen binding followed by sequencing to determine the distribution of variants across bins. **B**, Distribution of amino acid hamming distances of library members at different cycles compared to the parent sequence. **C**, A representative FACS plot showing the distribution of binding for the cycle 30 library and the thresholds used to sort cells based on relative binding. **D**, Normalized binding scores (see Methods) obtained from two independent biological replicates. Variants were classified into one of three classes (Parent-like, active, and lowly active) using a K-means clustering algorithm with K=3 and using normalized binding scores as features. **E**, Schematic overview of our approach to identifying matched sequences differing by a single amino acid mutation. In the example shown, the matched sequences differ by a glycine to aspartate mutation at the fourth position. **F**, The impact of mutation G61D across different variants. The colors for each variant correspond to the class assigned in panel E.

We next sought to profile the activity of individual sequences generated by the diversification campaign. To this end, we subcloned the diversified library onto a standard yeast nuclear plasmid, ensuring that each cell expresses and displays a single variant. Expression of the pooled library and incubation with IL-7Rα at the same permissive concentration used during diversification revealed a range of binding activities (Figure 3C). We then sorted cells based on their binding strength and used NGS to reconstruct the relative binding activity of each library member (Figure 3C). In total, we recovered binding data for 468 unique Mb-4 variants (Figure 3C and Figure S13). Of these, a majority (268/468) exhibited binding similar to the parental variant, and most variants (387/468) retained an appreciable binding phenotype. Together, these results demonstrate that OrthoRep can be harnessed to drive extensive diversification of small protein binders under permissive selection conditions to generate diverse sequence-affinity maps.

### Sequence context influences the impact of mutations on binding

The fitness effect of a mutation can depend on its surrounding sequence context^29,30^. The diverse set of variants generated through our neutral drift campaign provided a unique opportunity to explore how sequence context influences the contribution of specific mutations to binding. We first sought to determine the impact of individual mutations on the genetic background of the parental sequence by carrying out a deep mutational scanning (DMS) experiment on Mb-4. In this experiment, we generated a comprehensive library of Mb-4 variants containing single amino acid substitutions and characterized their binding relative to the parental sequence (Figures S14-S16). This allowed us to assess the functional impact of mutations at each position and calculate positional entropies—a measure of how tolerant each position is to mutation while still maintaining parent-like binding^11^. As expected, we observed low entropy in regions predicted to contact IL-7Rα, indicating strong conservation, and higher entropy at surface-exposed regions not predicted to directly participate to binding (Figure S14D), further supporting the validity of the predicted binding model (Figure S8). Moreover, the DMS dataset provides the effects of all single mutations for analysis against multi-mutant neutral drift outcomes.

To evaluate how mutational effects varied across different sequence backgrounds, we identified “matched” pairs of sequences from our neutral drift campaign that differed by only a single mutation (Figure 3E). This allowed us to isolate the effect of individual mutations across diverse mutational contexts. In some cases, the observed effects were consistent with those seen in the DMS data in the parental background (Figure S17A). However, other mutations exhibited context dependence (Figure S17B). A notable example was mutation G61D, located in a flexible linker region in Mb-4. While this mutation improved binding in the parental background (Figures S15 and S16), its effect in the neutral drift outcomes library varied widely—from nearly neutral to abolishing binding—depending on the surrounding sequence (Figure 3F). These results highlight that mutational effects are not fixed but can vary significantly depending on sequence context and that deep mutational scans in a single background may not fully capture the functional landscape of a protein as it evolves.

## Discussion

Recent breakthroughs in computational protein design have transformed our ability to engineer binders targeting proteins of interest including at user-specified epitopes^11–14,17,31,32^. Moving forward, two key challenges remain: improving hit rates (*i.e*., the percentage of designed binders that are functional) especially against challenging targets and reliably reaching high affinity. Toward these goals, we 1) demonstrate how OrthoRep-driven evolution can be harnessed to rapidly increase the binding affinity of computational minibinder designs and 2) systematically explore the mutational landscapes of affinity matured and neutrally drifted binder sequences, gaining insights into the complexities of design and datasets that may inform future design models.

In the first part of our work, we explored evolutionary outcomes when minibinders were subjected to stringent positive selection for improved binding. Despite their small size, minibinders responded robustly to stringent selection for improved binding, acquiring a handful of mutations that enhanced their affinity toward IL-7Rα. These mutations emerged over six to nine AHEAD cycles—corresponding to less than one month of experimental time (~3 days per cycle). This timescale is comparable to previous efforts from our lab to affinity mature nanobodies^12,24,25^. We used a high-affinity evolutionary outcome of one of the minibinders, Mb-1, to create a library of 64 mutants that represents a combinatorially complete sequence landscape for the six constituent mutations in the evolved Mb-1 outcome. The resulting sequence-affinity landscape, obtained using Tite-seq, is notable for two key reasons. First, they helped contextualize the contributions of mutations at residues involved in target binding versus those that are not. Specifically, the contributions of mutations not involved at the binding interface rival those of the mutations involved in binding. For example, the mutation that most improved binding, A14V, was not predicted to contact the target protein and may instead help stabilize the hydrophobic core of Mb-1. These findings, alongside prior work showing that *in silico* affinity maturation of antibodies is possible with general protein language models^33,34^—likely through stability-enhancing mutations—suggest that protein stability may be an important optimization criteria in computationally designed binders. Second, the binding landscape of these six affinity-improving mutations that connect the parental Mb-1 to its evolved outcome appears largely linear. Our hypothesis is that this reflects both the fact that we carried out affinity maturation through stringent selection, thus preferring a greedy search where single affinity-enhancing mutations successively fixed, and the simple structural topology of minibinder designs that present fewer opportunities for interactions among mutations.

In the second part of our work, we investigated the outcomes of minibinder evolution arising from a neutral drift process that prioritized the generation and maintenance of diverse sequences under purifying selection for function. We note two key outcomes of this neutral drift campaign. First, most of the final variants displayed binding comparable to the parental variant, whose activity defined our selection window (Figure 3A). The extensive diversification observed (8.3% divergence relative to the parental variant) suggests that even these small computationally designed proteins can tolerate substantial mutations while remaining functional and supports diversification from a single variant as a viable strategy to diverge a single binder into a large clade of similarly functional binders. Second, the results of the diversification campaign revealed sequence-dependent mutational effects, highlighting the importance of epistasis in shaping protein function. More broadly, this work underscores the importance of considering sequence context when interpreting mutational data or engineering protein variants. While single-mutation scans offer valuable insights, they may not fully capture the landscape of mutational effects once additional mutations accumulate. As such, approaches that retain or even encourage sequence diversity—like the permissive selection regime described here—can help reveal potential epistatic interactions that underly protein evolution, ultimately informing more robust strategies for protein engineering and design. Additionally, the variants that emerge from these diversification campaigns may serve as a useful resource when building and improving data-driven models for protein binder design, which we partially explored using a pre-trained language model^35^ (see Supplementary Text; Figures S18 and S19).

We envision several future adaptations that build on the approaches described here. First, while our evolution campaigns were each carried out in a single replicate, the scalability of OrthoRep should enable replicate evolution, facilitating the observation of reproducible or alternative evolutionary outcomes. Second, our neutral drift campaign began from only a single variant (*i.e*., the parental version of Mb-4). However, recent advances to OrthoRep enable evolution starting from large libraries^36,37^. For example, an error-prone PCR library could serve as a starting point to jump start extensive divergence. Third, while our present work focused on computationally designed minibinders, recent work introduced an approach to design single domain antibodies, such as nanobodies and single-chain variable fragments^12^, which were subsequently subjected to OrthoRep-driven evolution under strong selection. Instead of strong selection, the neutral drift approaches described in this work can also be applied to computationally designed antibodies in order to gain diverse sequence-affinity datasets that contain the epistatic complexities of sequence-binding landscapes to further train design models. Lastly, we could extend these approaches to evolve and study binders in contexts beyond yeast display—such as the intracellular recruitment of transcriptional or chromatin regulators^38,39^. Specifically, fusing a nanobody to a DNA-binding domain (*e.g*., zinc finger) could enable selection for transcriptional activator recruitment by linking binding to the expression of a prototrophic marker (for growth-based selection) or a fluorescent reporter (for FACS-based selection). Altogether, the convergence of *in silico* design and evolution offers a productive and mutually beneficial path forward for engineering protein binders and studying protein-protein interactions more broadly.

## Author contributions

M.A.A., C.E.T., C.J.L., I.C., and C.C.L. conceptualized the study. M.A.A., A.M.P., G.L.J.K., S.M., I.C.,

D.B., and C.C.L designed the experiments. M.A.A., A.M.P., and S.M. performed the experiments. D.B. led the deep mutational scanning experiments with help from S.J.H. and G.C. M.A.A., A.M.P., S.M., D.B., G.L.J.K., I.C., and C.C.L. analyzed the data. H.B., S.M., and S.J.H. selected minibinders 1-4 for this study from a *de novo* design/yeast display discovery campaign executed by H.B., S.J.H., G.C, and S.M. under the supervision of C.E.T. and I.C. M.A.A. wrote the software to analyze the Tite-seq and neutral drift library sequencing data. H.B. and G.K.J.K. performed structural modeling using AF2. T.S., M.J., B.J., and S.V.I. performed machine learning modeling. G.L.J.K. generated Figure S14D, and T.S. generated Figure S18. M.A.A. and C.C.L. created the visualizations for all other Figures. C.E.T., C.J.L., I.C., and C.C.L. supervised the study. M.A.A. and C.C.L. wrote the paper with input from all authors.

## Acknowledgements

We thank Derek Aspacio, Jacob Hambalek, Vincent Hu, Rory Williams, Yutong Yu, and the members of the Liu Lab for their advice and support; Lydia Green and Claire Liu for assistance with minibinder production; Desiree Lim for minibinder construct design and cloning and reagent preparation; Lindsey Gross, Nina Takahashi, Dominik Neef, Yoan Machado Hernandez, and Nastaran “Nancy” Nosrat Tarjoddin for reagent preparation; Bhuvaneswari Kasi, Chetan Lingaraju, S.P. Prashanth, Janak Prasad and Daniel Lu for their assistance with deep mutational scanning; Robert Vernon for discussions on the use of AMPLIFY. This work was funded by a sponsored project from Amgen to C.C.L. M.A.A. was supported by a University of California Chancellor’s Postdoctoral Fellowship. We thank the Institute for Rapid Antibody Engineering and Evolution, part of the Engineering+Health Initiative of the UCI Samueli School of Engineering, and a research grant from the W.M. Keck Foundation to C.C.L. for additional support.

## Data availability

The raw sequencing files will be made publicly available under NCBI Bioproject PRJNA1304565.

## Code availability

All code for analyzing the combinatorial Mb-1 library and diversified Mb-4 library will be made available on GitHub (https://github.com/maalcantar/minibinders_orthorep_analysis).

## Materials and Methods

### Strains, media, and routine plasmid construction

Key strains and plasmids are described in Tables S4 and S5. All routine cloning was performed using NEB Stable (New England Biolabs) or One Shot™ TOP10 (Thermo Fisher Scientific) *Escherichia coli* competent cells. *E. col*i were grown at 37 °C in 2-YT Broth consisting of 16 g/L Tryptone (US Biological), 10 g/L yeast extract (US Biological), and 5 g/L NaCl (US Biological) with the appropriate antibiotic selection for plasmid selection (100 μg/mL ampicillin/carbenicillin or 50 μg/mL kanamycin). Plasmids were constructed using Gibson Assembly and Golden Gate Assembly.

The background strains used for all experiments in this study were the previously published strains yAP174 and yAP196 (ref^25^), which were derived from *Saccharomyces cerevisiae* BJ5465 (MATa ura3-52 trp1 leu2-delta1 his3-delta200 pep4::HIS3 prb1-delta1.6R can1 GAL). Briefly, yAP174 was used to express AGA2-tethered minibinder proteins from a nuclear CEN/ARS plasmid and contains three changes relative to *S. cerevisiae* BJ5465. First, the native *AGA1* locus was replaced with an *AGA1* gene controlled by a β-estradiol inducible promoter (pER; a CYC1 minimal promoter with upstream zinc finger [ZF] binding sites for ZF 97-4). Second, the native *URA3* locus was replaced with a synthetic transcription factor (synTF; a fusion protein consisting of ZF 97-4, estradiol-responsive nuclear localization domain, and VP16 minimal activation domain) which induces expression from pER in the presence of β-estradiol^40^. Lastly, a complete deletion of the native *trp1* gene was incorporated, resulting in yAP174. Three additional modifications were made to yAP174 to create yAP196, which was the strain used to affinity mature and diversify the minibinder proteins. First, a complete deletion of the native *MET15* gene was incorporated. Second, a landing pad p1 plasmid and p2 plasmid expressing genes required for linear plasmid replication were added through a protoplast fusion using an F102-2 strain as the donor. Lastly, a CEN/ARS plasmid harboring the BadBoy3 error-prone polymerase was transformed^23^, resulting in yAP196.

All yeast strains in this study were cultured at 30 °C with shaking at 200 rpm in synthetic complete (SC) growth medium: 6.7 g/L yeast nitrogen base without amino acids and with ammonium sulfate (US Biological), 20 g/L dextrose (Fisher Scientific), and the appropriate nutrient drop-out mixture at the concentration recommended by the manufacturer (US Biological). In cases where a eukaryotic antibiotic (*e.g*., nourseothricin) was used as part of the selection, the yeast nitrogen base was replaced with 1.72 g/L yeast nitrogen base w/o amino acids w/o ammonium sulfate (US Biological) and 1 g/L of L-Glutamic acid monosodium salt hydrate (Sigma-Aldrich). To prevent bacterial contamination of yeast cultures, 50 μg/mL carbenicillin and 15 μg/mL chloramphenicol were added to the synthetic media.

Lastly, all references to IL-7Rα refer to the human homolog of the receptor (Table S1; GenBank accession: AAC83204) unless otherwise specified.

### Yeast transformations

Yeast cells were made competent following the protocol described by Gietz and Schiestl^41^ unless otherwise specified. In brief, a single yeast colony was picked into 5 mL of the appropriate media and grown at 30 °C to saturation. From the saturated culture, 3.3 mL was transferred to 100 mL of YPD media (20 g/L peptone, 10 g/L yeast extract, 20 g/L dextrose) and grown until the optical density at 600 nm (OD_600_) was between 0.6 and 1.0 (~6-8 h). Once the appropriate OD_600_ was reached, 100 mL of the culture was transferred to two 50 mL conical tubes and pelleted at 3,000 × g for 5 min at room temperature. The supernatant was discarded, and each pellet was washed with 25 mL of sterile water. Next, the cells were pelleted again at 3,000 × g for 5 min, the supernatant discarded, and the cells were resuspended in 500 μL of sterile water. The resuspended cells were transferred to a 1.5 mL microcentrifuge tube and pelleted at 3,000 × g for 5 min. The supernatant was removed, and the cells were resuspended in 500 μL of frozen competent cell storage solution consisting of 5% v/v glycerol and 10% DMSO in sterile water. 50 μL aliquots of the resulting competent cells were either stored at −80 °C or used immediately for transformations.

For yeast transformations, the competent cells were thawed (if previously frozen), pelleted at 12,000 × g for 2 min, and the supernatant was discarded. The pelleted cells were resuspended in 260 μL of 50% poly(ethylene glycol), 36 μL of 1 M lithium acetate, 20 μL of pre-boiled salmon sperm DNA (5 mg/mL), purified DNA construct, and nuclease-free water. The resuspended cells were incubated at 42 °C for 20 min. Following this incubation, the cells were pelleted at 13,000 × g for 30 s and the supernatant was removed. The cells were resuspended in 1 mL of YPD and incubated at 30 °C for either 1 h (for prototrophic selections) or 6-8 h (for antibiotic selections). The cells were then pelleted at 13,000 × g for 30 s, washed with 1 mL 0.9% NaCl, and pelleted again. The resulting pellet was then resuspended in 500 μL 0.9% NaCl and plated on selective media.

For transformations to install new constructs onto the p1 landing pad in strain yAP196, 3 μg of a donor plasmid (pMAA27) containing homology to the landing pad was digested in a 50 μL reaction containing 2 μL of ScaI-HF (New England Biolabs) for at least 3 h at 37 °C followed by a heat inactivation at 80 °C for 20 min. Complete digestion was confirmed by running 5 μL of the reaction on a 0.9% TAE agarose gel. The remaining 45 μL of digest product was added to the transformation mixture described above.

### Computational design of the four minibinders and initial screening by yeast display

Minibinder sequences were designed using an approach similar to that described by Cao et al^11^. Briefly, a scaffold library of trihelical bundles and ferredoxin folds were constructed using the blueprint builder^42^ in RosettaRemodel^43^. The IL-7 binding site was selected as the IL-7Rα epitope for design. IL-7Rα residues having ≥10 Å^2^ difference in solvent accessible surface area between IL-7-bound and free states, based on the IL-7Rα crystal structure (PDB ID: pdb_00003di3; ref.^44^), were used to generate a rotamer interaction field (RIF). This RIF was used to dock scaffolds to IL-7Rα which was followed by a round of ProteinMPNN^19^ sequence design. AlphaFold-multimer^21^ was used to filter out designs having pLDDT < 90 or backbone atom RMSD from design ≥ 1 Å. Mb-4 was further refined in an additional round of Rosetta FastDesign restricted to amino acids more than 12 Å from the interface. Disulfide bonds were introduced into Mb-1 and Mb-2 using the stapler PyRosetta package^45^.

The designed minibinder sequences were codon optimized for *S. cerevisiae*, inserted between invariable 5′ and 3′ primer binding sites to enable amplification, and synthesized as an oligonucleotide pool by Twist Bioscience. A library DNA fragment with 5′ and 3′ overhang sequences was generated via PCR amplification of the pool to support homologous recombination with a yeast display vector encoding a fusion protein consisting of Aga2p-hemagglutinin (HA) tag-minibinder coding sequence-Myc tag under a galactose-inducible promoter, as described in Chao *et al*^46^. The linearized plasmid and DNA fragment for the computationally designed minibinder library were transformed into EBY100 yeast strain as described by Benatuil *et al*^47^. Minibinder sequences were displayed and labeled for fluorescence-activated cell sorting (FACS). Target binding at 50 nM IL-7Rα-Fc-Biotin (AcroBio; catalog #IL7-H82F8) was assessed using a streptavidin-phycoerythrin conjugate (ThermoFisher; catalog #S866), and minibinder surface expression was inferred by staining with an anti-HA-AlexaFluor 647 conjugate (ThermoFisher; catalog #26183-A647). After FACS-based stratification, a subset of sequences was identified for recombinant expression and purification.

### Affinity maturation of the four minibinders

The minibinders used in this study were cloned onto a p1 donor plasmid (pMAA27) using Gibson Assembly. This resulted in a gene encoding an in-frame fusion protein consisting of an app8 signal peptide, minibinder, hemagglutinin (HA) peptide tag, and AGA2 controlled by pGA which is a p1-specific promoter. These donor constructs were integrated onto the p1 landing pad in yAP196 as described above, and transformants were selected by plating on SC-HLUW with nourseothricin (100 μg/mL). Nourseothricin was only used during this initial selection to enrich for colonies that have undergone successful integration onto p1 and prevent the formation of “cheater” colonies that fulfill their tryptophan auxotrophy by feeding off neighboring cells that have undergone a successful integration.

Individual colonies were inoculated into SC-HLUW media and grown to saturation. Once cells reached saturation, they were passaged 1:100 into SC-HLUW with 100 nM β-estradiol to induce display of the minibinder and grown overnight (16-24 h).

Approximately 5×10^7^ cells were taken from the induced culture and used to measure binding to human IL-7Rα. First, the cells were pelleted at 3,000 × g for 5 min and washed twice with 250 μL of sorting buffer (20 mM HEPES-NaOH at pH 7.5, 100 mM NaCl, 5 mM maltose, and 0.1% BSA). The washed cell pellet was resuspended in 250 μL of the sorting buffer containing purified human IL-7Rα. For most experiments, an IL-7Rα variant fused to a human IgG single-chain Fc tag (synthesized and purified in-house) was used. In a few cases, a biotinylated version of IL-7Rα (Acro Biosciences) was used. The resuspended pellet was incubated at 4 °C for 1 h with rotation to allow for IL-7Rα binding. The cells were then pelleted at 3,000 × g for 1 min and washed twice with 250 μL of the sorting buffer. The cell pellet was resuspended in 250 μL of sorting buffer containing 2.5 μL of anti-HA-AF488 (R&D systems; catalog #IC6875G-100UG) and 0.5 μL of anti-human IgG-AF647 (Southern Biotech; catalog #9040-31) for the Fc-tagged IL-7Rα or streptavidin-APC (Thermo Fisher; catalog #17-4317-82) for the biotinylated IL-7Rα. This mixture was incubated at 4 °C for 30 min with rotation. Next, the cells were stained with 5 μL of propidium iodide (1 mg/mL stock solution) incubated for 1 min at 4 °C, then pelleted at 3,000 × g for 1 min and washed twice with sorting buffer. The pellet was then resuspended in 4 mL of sorting buffer in preparation for sorting. FACS was performed (Sony SH800) and cells were sorted into 3 mL of SC-HLUW. Approximately 250-1,000 cells were sorted during each round of selection. The sorted cells were grown to saturation and the process of induction, staining, and selection was repeated. The selection gates and concentrations used during each round of affinity maturation are shown in Figures S2-S5. The p1 sequences from the final cycles were amplified using PCR and cloned into a CEN/ARS display library. For each minibinder, two final sorting steps were performed to purify clones containing the best binders. After the second sort, cells were plated, and individual colonies were sequenced and tested for binding. One representative binder from each minibinder evolution is reported in this study.

### On yeast dose-response curves

The minibinders used in this study were cloned onto a CEN/ARS plasmid for yeast display (pMAA28). A single yeast colony was picked into selective media (SC-HLU) and grown overnight to saturation. The saturated cultures were then passaged 1:100 into selective media containing 100 nM β-estradiol and grown for 16 h to induce minibinder expression. Cultures were pelleted at 750 × g for 5 minutes, washed twice with sorting buffer, and resuspended in 1 mL of sorting buffer. An aliquot of the resuspended cells was diluted into sterile sorting buffer which was used to obtain accurate cell densities by using an Attune flow cytometer (Life Technologies) to measure cell counts. Once cell counts were obtained, cells were diluted into 5 mL of sorting buffer at a density of 10^6^ cells per mL. In a 96-well V-bottom plate, fourfold serial dilutions of IgG-Fc tagged IL-7Rα starting from 100 nM (2× working concentration) were prepared in 100 μL volumes. To these serial dilutions, 100 μL of the yeast suspension (approximately 1×10^5^ cells) were added. The cells were then incubated at 4 °C with shaking at 200 rpm for 1 h. After this incubation, the cells were pelleted at 2,800 × g for 5 minutes and washed twice with sorting buffer. The cell pellet was resuspended in 50 μL of a secondary staining solution consisting of sorting buffer with 1 μL of anti-HA-AF488 and 1 μL of goat anti-human IgG-PE (Abcam; catalog #ab131612). The cells were incubated at 4 °C (200 rpm) for 30 min, pelleted, washed twice, and then resuspended in 150 μL of sorting buffer. Display and binding were measured on an Attune flow cytometer (Life Technologies). To quantify binding, the median fluorescence intensity of binding (*i.e*., PE signal) was computed for the displaying population.

### AlphaFold3 predictions and PyMOL visualization

Structural predictions for the minibinders and human IL-7Rα were generated by UC Irvine using AlphaFold3 (ref^26^), accessed via the AlphaFold Server. Predictions were performed using the amino acid sequences of the parent minibinders and the recombinant extracellular domain fragment of IL-7Rα that was used in this study (Table S1). For all predictions, the random seed was set to ‘42’ to ensure reproducibility. The predicted structures were visualized and colored in PyMOL (The PyMOL Molecular Graphics System, Version 3.0.2, Schrödinger, LLC).

### Dissociation constant measurements (K_D_) using surface plasmon resonance (SPR)

Minibinder coding sequences were synthesized as linear expression cassettes containing a T7 promoter and terminator (IDT) and an N-terminal polyhistidine tag for affinity purification. Proteins were expressed in a cell-free format using the Juice Cell-Free Kit (Liberum Catalog CF0030.B) according to the product manual, in a T100 thermal cycler (BioRad). Minibinders were purified in a single immobilized metal affinity chromatography step using a KingFisher Apex (Thermo Scientific) and AmMag Ni magnetic beads (GenScript). Cell-free production of minibinders was confirmed by 4–20% Mini-PROTEAN TGX stain-free protein gel (BioRad). DNA encoding the human IL-7Rα extracellular domain (residues 37-231; Table S1) with a C-terminal C-tag was cloned into a mammalian stable expression vector and expressed in CHO-K1 cells^48^. Conditioned media was purified with a 5 mL CaptureSelect C-tagXL affinity column (Thermo Scientific) followed by polishing via size-exclusion chromatography on a 320 mL HiLoad 26/60 Superdex 200 pg column (Cytiva).

Binding kinetics were determined using a Carterra LSA high-throughput SPR instrument. Purified minibinders were first captured with a NiHC200M sensor chip to reach an average density of 200 RU, using a running buffer containing 0.1 M HEPES 1.5 M NaCl 0.05% v/v Tween 20, pH 7.4. Next, seven identical cycles of running buffer injections were performed to stabilize the NiHC200M biosensor and to remove any unbound material. The analyte, IL7-Rα extracellular domain (residues 37-231) with a C-terminal C-tag, was formulated to 1000 nM in running buffer and serially diluted 1:3 down to 4.11 nM Analyte was injected for 5 minutes at increasing concentrations, and each injection was followed by a 15-minute dissociation time. Data was double referenced using the standard Carterra spot referencing scheme and nearest blank subtraction. Kinetic constants were fit using a 1:1 Langmuir binding model using Kinetics 1.9.2.4430 software (Carterra) to capture Rmax, *k*a, *k*d and K_D_ values, with floated t_0_ for the non-regenerative kinetic mode and bulk shift set to constant 0.

### Mb-1 library construction

The combinatorially complete Mb-1 library was constructed by ligating oligonucleotide duplexes into a CEN/ARS yeast display backbone (pMAA28). A complete list of oligonucleotides used is provided in Table S6. The Mb-1 sequence was divided into four segments, each containing one to three nucleotide positions that were mutated in the affinity matured variant. For each segment, we synthesized 2^N^ oligonucleotide pairs where N is the total number of mutations in a given segment. Upon annealing, cognate oligonucleotides formed 4 bp overhangs that were unique to each segment, enabling ligation between neighboring segments and the backbone. Each minibinder variant was assembled by ligating the appropriate oligonucleotide duplexes with the linearized yeast display backbone (pMAA28; linearized with BsaI-HF). The assembled plasmids were then individually transformed into competent NEB Stable (New England Biolabs) *E. coli* cells and purified by miniprep. Plasmids encoding all 64 variants were pooled at equimolar ratios to create the combinatorially complete library.

### Tite-seq pipeline and NGS preparation

The pooled library was transformed into the yeast display strain yAP174 as described above, with the exception that transformed cells were directly inoculated into 50 mL of selective media. Biological replicates were obtained by retransforming the pooled library. The cells were grown to saturation and 300 μL of the saturated culture was added to 30 mL of selective media containing 100 nM β-estradiol and grown for 16 h to induce expression of the minibinder library. Aliquots containing 2 × 10^7^ cells from the induced culture were then stained and labeled as described above at seven concentrations of Fc-tagged IL-7Rα: 0, 0.01, 0.1, 1, 10, 100, and 500 nM.

At each concentration, FACS was performed (Sony SH800) to sort the cells into four bins based on IL-7Rα binding (*i.e*., AF-647 intensity). Prior to sorting, approximately one million events from cells stained with 0 and 500 nM IL-7Rα were measured to obtain the dynamic range of AF-647 intensity and establish sorting bins. The first bin captured all AF-647-negative cells, while the remaining bins were spaced equally apart in log-space, covering the full range of AF-647 intensity (Figure S9). For each IL-7Rα concentration, cells were collected into 3.3 mL of selective media in two batches. The first batch collected cells into bins 1 (no binding bin) and 3. The second batch collected cells into bins 2 and 4 (highest binding bin). In each batch, 2 million events were processed to normalize the number of cells sorted in each batch and across concentrations.

After sorting, the cells were grown to saturation, and 1.5 mL aliquots were used for minipreps to extract CEN/ARS plasmids harboring the minibinder variants. Briefly, 1.5 mL of saturated culture were pelleted at 8,000 × g, washed with 500 μL of 0.9% NaCl, and pelleted again. The cells were then resuspended in 250 μL of a Zymolase solution (0.9 M sorbitol, 0.1 M EDTA [pH 7.5], and 0.5 mg/mL Zymolase) and incubated at 37 °C for 1 h with rotation. After incubation, the cells were pelleted at 16,000 × g for 1 min, the supernatant was discarded, and the pellet was resuspended in 561 μL of a Proteinase K solution (500 ul TE [50 mM Tris-Cl at pH 7.5 and 20 mM EDTA], 50 μL of 10% SDS, and 11 μL of 10 mg /mL proteinase K). The mixture was incubated at 65 °C for 30 min, followed by the addition of 200 μL of 5 M potassium acetate and incubation at 4 °C for 30 min. The mixture was then centrifuged at 16,000 × g for 1 min, and the supernatant was column purified (EconoSpin® Mini Spin Column; Epoch Life Sciences) to yield purified plasmid DNA.

The purified plasmid DNA was used as a template for PCR amplification using primers that annealed to the *app8* signal peptide sequence and glycine-serine linker flanking the minibinder sequence. Bin-specific barcodes were incorporated into the PCR primers to allow pooling of amplicons within each IL-7Rα concentration. These barcodes were generated with the DNABarcodes library^49^ v1.28.0 in R v3.6.1, using the “create.dnabarcodes” function with parameters “n = 4 dist = 4 heuristic = ‘ashlock’”. This generated 4 bp barcodes that all possessed a Hamming distance of 4 bp with all other barcodes. The minibinder amplification was performed in a 50 μL PCR reaction containing 2 μL of the miniprep product, 5 μL of each oligonucleotide pair (*e.g*., MAA_o256 and MAA_o257 at 10 μM for miniprepped plasmid library from bin 1), and 25 μL of NEBNext Ultra II Q5 Master Mix. The PCR conditions were as follows: 98 °C 30 s → (98 °C 10 s → 65 °C 75 s) × 21 cycles → 65 °C 5 min → 4 °C hold. Following PCR, amplicons were column purified (EconoSpin® Mini Spin Column; Epoch Life Sciences). DNA concentration was quantified using a Qubit3 fluorometer (Thermo Fisher) with 2 μL of DNA. Additionally, 1.5 μL of the purified DNA was run on a 2.0% TAE agarose gel to confirm the expected amplicon size. For each IL-7Rα concentration, purified amplicons from the four bins were pooled together (resulting in seven pools per biological replicate) and submitted for paired-end sequencing via Azenta Life Sciences’ Amplicon-EZ service (2 × 250 cycles).

### NGS data processing and Tite-seq analysis

Raw paired-end reads were first demultiplexed using a custom Python script, which identified the bin-specific barcodes added during the minibinder PCR. The demultiplexed paired-end reads were then quality filtered and trimmed using fastp^50^ v0.23.2 with parameters “-l 150 -q 30 -u 20”, which enforced a minimal read length of 150 nucleotides and discarded reads where more than 20% of bases had a quality score below 30. The forward and reverse quality filtered reads were then merged using PEAR^51^ v0.9.11 with parameters “-n 238 -m 258 -j 7”. The filtered, merged reads were then mapped to the parent minibinder sequence using Bowtie2 (ref^52^) v2.4.4 with parameters “--sensitive-local --no-unal -p 7”. The mapped sequences were then extracted from the resulting BAM files and compared to the parent sequence to identify mutations. A count table of minibinder variants across concentrations and bins was then generated using a custom Python script.

The read counts of each minibinder variant across bins were used to infer mean fluorescence values for each IL-7Rα concentration. First, normalized fractional abundances 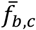 for a given minibinder variant in bin *b* at a given IL-7Rα concentration *c* was computed as:

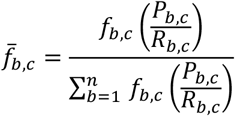

where *f*_*b,c*_ was the raw read count for a given minibinder variant in bin *b, P*_*b,c*_ was the proportion of cells sorted into bin *b, R*_*b,c*_ was the total number of sequencing reads obtained from bin *b*, all at a given IL-7Rα concentration *c*. Also, *n* was the number of bins used (*n*=4 in our experiments). These normalized fractional abundances were used to compute mean fluorescence 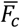 at a given IL-7Rα concentration using:

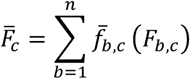

where *F*_*b,c*_ is the geometric-mean of fluorescence for all cells in bin *b* at IL-7Rα concentration *c*. K_D_ values were then inferred by fitting the mean fluorescence values to a Hill function of the form:

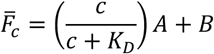

where *A* is the fluorescence increase at saturating IL-7Rα concentrations and *B* is the background fluorescence in the absence of IL-7Rα. Curve fitting was implemented in Python v3.8.19 using SciPy^53^ v1.10.1 with the sp.optimize.curve_fit function. The following boundary conditions were applied: *K*_*D*_ [10^−5^, 10^5^], A [0, 10^6^], and B [0,10^5^]. Fits were computed for each biological replicate, and the reported K_D_ values represent the mean from two biological replicates. The dose-response curves shown in Figure S10 depict log_10_-transformed 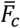 values. The combinatorial binding landscape shown in Figure 2B was created using NetworkX^54^ v3.1 with −log_10_(K_D_) values, where higher values indicate stronger affinity.

### Modeling the effects of mutations on binding

All models were implemented using scikit-learn^55^ v1.3.2 in Python v3.8.19. For all models, training and testing were performed using an 80/20% data split with 100 different train-test splits to ensure model robustness. We implemented a linear model to infer the effects of individual mutations, as previously described^30,56^. Briefly, the model assumes that −log_10_(K_D_) for a given minibinder variant *m* can be described as a sum of individual mutational effects as given by:

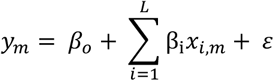

where *y*_*m*_ is the −log_10_(K_D_) for minibinder variant *m, β*_*o*_ is the −log_10_(K_D_) for the parental variant, β_i_ represents the effect of mutation *i* on binding, *x*_*i,m*_ indicates whether mutation *i* is present in minibinder variant *m* (1 if present, 0 otherwise), and ε represents the error term. The model was implemented with L2-regularization using the sklearn.linear_model.Ridge function with parameters “α = 0.0001, solver = ‘lsqr’, fit_intercept = False”.

Higher-order models, which include epistatic interactions, were implemented using the following framework:

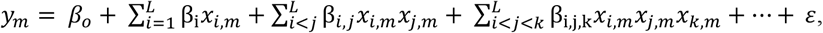

where, β_*i,j*_ represent second-order interaction coefficients between mutations *i* and *j*, β_*i,j,k*_ represents third-order interaction coefficients between mutations *i, j*, and *k*. The same form can be extended to capture higher-order interactions. The reported coefficients for the models are the mean across 100 train-test splits.

### Deep mutational scanning of Mb-4

An SSM (single-site saturation mutagenesis) library was designed to incorporate randomization across the entire minibinder sequence using the NNK codon (N = A/T/G/C, K = G/T). The DNA fragments encoding the SSM library were codon optimized for *S. cerevisiae* using tools on the TelesisBio Website (https://telesisbio.com/). Each Mb-4 library DNA sequence was flanked by invariant 5′ and 3′ overhang sequences to enable homologous recombination as described above for initial yeast display screening. The library fragments were synthesized and assembled using the BioXp 9600 system (Telesis Bio) and pooled prior to transformation into EBY100 yeast. For FACS, target binding was assessed at 10 nM of single-chain-Fc-tagged human IL7Ra-Single Chain Fc (ScFc) followed by detection with anti-huIgGFc-Phycoerythrin (PE) antibody (Invitrogen 12-4998-82). Surface display was assessed by anti-HAtag-AF 647 (Invitrogen 26183-A647). Cell sorting was carried out using a BD-FACS Melody sorter. Cells were first gated on single cells and then stratified to yield four different binding bins (high, medium (parent-like), low, and non-binding) based on their PE fluorophore signal^57^. A parallel sort collecting the entire HA tag-positive population served as the expression-sorted reference population.

After one round of FACS, plasmid DNA was extracted from all sorted yeast pools using the Zymoprep yeast plasmid miniprep II kit (D2004). The entire minibinder sequence was amplified from the isolated yeast plasmid prep with vector-specific primers and purified using the Qiagen gel-extraction kit. Sequencing libraries were constructed from purified amplicons using Illumina NexteraXT PCR (Catalog no. FC-131-1096) and sequenced on the Illumina NextSeq2000 platform with a 2×300 paired-end read configuration and dual 8 base pair indexes. NGS data analysis was performed using proprietary software following the same principles are previously described^58^. Briefly, FASTQ files from forward and reverse reads were merged using NGmerge^59^ v0.4 and the concatenated reads were further trimmed to yield only the minibinder sequence for downstream analysis. Over 3×10^6^ minibinder reads from each sort pool were analyzed as follows. Minibinder sequences were aligned with the reference parent sequence to identify mutations versus the parent and generate read counts. To ensure data robustness and minimize noise from low-abundance variants, mutations with fewer than 100 counts in the expression sorted pool were excluded and denoted “not captured.” Variants which were absent in binding-sorted pools but present in the expression-sorted pool were assigned a read pseudocount of 0.5 (rather than 0) to enable valid calculations of binary logarithms [*i.e*., log_2_(enrichment factor)]. Binding enrichment factors (EF) for each mutation were calculated by comparing frequency in each binding bin to the reference population (EF = freq_bind pool_/freq_reference (expression) pool_), converted to binary logarithms, and normalized to that of the parent minibinders as previously described^58^.

### Using OrthoRep to diversify Mb-4 through neutral drift

Yeast strains containing Mb-4 in the p1 plasmid were inoculated from glycerol stocks into selective media and grown to saturation. The saturated culture was then diluted 1:30 into selective media containing 100 nM β-estradiol and grown for 16 h. From the induced culture, 5 × 10^7^ cells were labeled and stained as described above using 25 nM of IL-7Rα. In the initial sort, a gate was drawn to encompass the founder population, selecting cells within this binding and display regime. The gate was slightly adjusted in subsequent sorts to account for variability, and the final sorts included a broader gate to capture a more diverse range of minibinder variants. The sorted cells were collected into 4 mL of selective media and grown to saturation. The process of passaging, induction, and sorting was repeated for a total of 30 cycles. During the initial cycle, approximately 10,000 cells were sorted. In subsequent cycles, approximately 80,000 cells were sorted per cycle. The sorting gates used during each cycle are shown in Figure S15.

After reaching saturation, the sorted cultures from cycles 8, 15, and 30 were miniprepped, and the purified p1 plasmids were amplified in preparation for NGS as described above. Amplification was performed using primers K221 and K222 (Table S6). The purified amplicons were sequenced using Azenta Life Sciences’ Amplicon-EZ service (2 × 250 cycles). Minibinder variants were identified by mapping reads to the parent sequence using Bowtie2 (ref^52^). Only unique amino acid variants with at least 10 reads were included in Figure 3B.

### Building the Mb-4 neutral drift outcome library

The diversified Mb-4 library, post-cycle 30, was cloned into a CEN/ARS backbone (pMAA28) using Gibson Assembly in preparation for library characterization. Briefly, 1.5 mL of saturated culture from the post-cycle 30 sort was miniprepped as described above. The minibinder variants were amplified from the purified p1 plasmid in a 50 μL PCR reaction containing 2 μL of miniprep product, 2.5 μL of each oligonucleotide pair (*i.e*., MAA_o254 and MAA_o255 at 10 μM), 10 μL Q5^®^ reaction buffer, 1 μL of dNTPs at 10 mM, and 0.5 μL of Q5^®^ High-Fidelity DNA Polymerase. The PCR conditions were as follows: 98 °C 30 s → (98 °C 10 s → 65 °C 75 s) × 27 cycles → 72 °C 2 min → 4 °C hold. The resulting amplicons were column purified, quantified on a Qubit3 fluorometer (Thermo Fisher), and verified on a 2.0% TAE agarose gel. The CEN/ARS yeast display backbone (pMAA28) was linearized in a 40 μL reaction containing 3 μg of plasmid, 2 μL of BsaI-HF (New England Biolabs), and 4 μL of CutSmart^®^ buffer (New England Biolabs). The reaction was incubated for 12 h at 37 °C followed by a heat inactivation at 80 °C for 20 min. The entire digest reaction was run on a 0.9% TAE-agarose gel and the backbone was gel extracted. The minibinder amplicons and linearized backbone were assembled in 12 μL Gibson Assembly reaction at 50 °C for 1 h. The Gibson Assembly reaction then underwent ethanol precipitation to obtain purified DNA. The ethanol precipitation was performed as follows. First, 38 μL of nuclease-free water was added to the Gibson reaction, bringing the total volume to 50 μL in a 1.5 mL microcentrifuge tube. Then, 5 μL of sodium acetate (pH 5.2) and 1 μL of glycoblue (Thermo Fisher) was added to aid in visualization of the DNA pellet in subsequent steps. To this mixture, 125 μL of ice-cold 100% ethanol was added, and the mixture incubated at −80 °C for 20 min. After the incubation, the mixture was centrifuged at 12,000 × g for 30 min at 4 °C. The supernatant was removed, and the pellet was washed with 175 μL of ice-cold 70% ethanol. The mixture was centrifuged for 10 min at 4 °C, the supernatant was removed, and residual ethanol was evaporated by air-drying for 10 min at room temperature. The DNA pellet was then resuspended in 10 μL of nuclease-free water and 3 μL of the resulting pool of plasmids was used to transform electrocompetent One Shot™ TOP10 *E. coli*. After the electroporation, cells were recovered in SOC media for 1 h. Recovered cells were inoculated directly into 2-YT with carbenicillin (100 μg/mL) and an aliquot was serially diluted and plated on 2-YT agar plates with carbenicillin to assess transformation efficiencies. Colony counts confirmed that transformation efficiencies exceeding 10^6^ were achieved. The transformed cells grown in 2-YT with carbenicillin were grown to saturation and miniprepped to obtain purified plasmid library.

### Sorting the Mb-4 neutral drift outcome library, NGS preparation, and analysis

The purified Mb-4 plasmid library was transformed into the yeast display strain (yAP174) and induced as described above: grown to saturation, passaged 1:100 into selective media with 100 nM β-estradiol, and grown for 16 h. Alongside the library, cells were also independently transformed with the parental Mb-4 to help establish the parental binding population in subsequent sorting. After induction, 2 × 10^7^ cells were stained and labeled with 25 nM IL-7Rα and labeled as described above. The isogenic population expressing the parental minibinder was first measured to help establish bins based on binding levels. Bin 1 was adjusted to capture the binding-negative (AF-647 negative) populations. Bins 2, 3, and 4 were adjusted to capture cells with lower, similar, and higher binding, respectively, compared to the parental variant. Once the gate thresholds were established, the library was sorted in two batches. The first batch collected cells into bins 1 (no binding) and 3 (medium binding). The second batch collected cells into bins 2 (low binding) and 4 (high binding). For each batch, 8 million events were processed to normalize the number of cells sorted in each batch. The cells were collected into 4 mL of selective media and grown to saturation. From the saturated cultures, 1.5 mL aliquots were miniprepped to purify plasmids. The purified plasmids were used as templates for PCR in 50 μL reactions containing 2 μL of miniprepped DNA, 5 μL of the forward and reverse primer (K221 and K222 at 10 μM), and 25 μL of NEBNext Ultra II Q5 Master Mix. The PCR conditions were as follows: 98 °C 30 s → (98 °C 10 s → 65 °C 75 s) × 21 cycles → 65 °C 5 min → 4 °C hold. The purified amplicons were quantified on a Qubit3 fluorometer (Thermo Fisher) and verified on a 2% TAE agarose gel. Purified amplicons submitted for paired-end sequencing through Azenta Life Sciences’ Amplicon-EZ service (2×250 cycles).

Raw, paired-end reads were quality filtered, trimmed, merged, and mapped to the parent sequence as described above. The only difference was that PEAR^51^ v0.9.11 was used with parameters “-n 268 -m 288 -j 7”. All minibinder variants were collapsed at the amino acid level, meaning variants with different DNA encodings for the same amino acid sequence were aggregated. Only amino acid variants with more than 20 reads across all bins in both biological replicates were used in downstream analyses.

Normalized binding scores were computed using a slightly modified approach from what was previously used to process the Tite-seq data. First, normalized fractional abundances 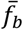for a given minibinder variant in bin *b* was computed as:

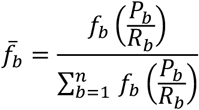

where *f*_*b*_ was the raw read count for a given minibinder variant in bin *b, P*_*b*_ was the proportion of cells sorted into bin *b, R*_*b*_ was the total number of sequencing reads obtained from bin *b*, and *n* was the number of bins used (n=4 in our experiments). Binding scores were then scaled between [0,1] by taking a weighted averaged of the fractional abundances for minibinder variants across each bin:

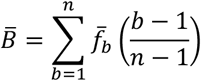

where 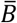was the normalized binding score for a given minibinder variant. A small number of sequences where replicate normalized binding scores differed by more than 0.33 a.u., roughly translating to the replicates being more than one bin apart, were discarded.

The minibinder variants were classified into one of three classes (lowly active, active, and parent-like) with a K-mean clustering algorithm using the normalized binding scores for both biological replicates as features. The K-means clustering algorithm was implemented in scikit-learn^55^ v1.3.2 in Python v3.8.19 using the KMeans function with parameters “n_clusters = 3, random_state = 42, n_init='auto'“.

Two-dimensional representations of the minibinder variants were computed with the pacmap^60^ v0.7.6 package using the PaCMAP function with parameters “n_components = 2, n_neighbors = 10, MN_ratio = 0.5, FP_ratio = 2.0, random_state = 42.” The two-dimensional representations were also clustered using the KMean function with parameters “n_clusters = 4, random_state = 42, n_init='auto'“. Sequence logos were generated using the logomaker^61^ v0.8.6 package.

## References

1. Dueber, J. E., Yeh, B. J., Chak, K. & Lim, W. A. Reprogramming Control of an Allosteric Signaling Switch Through Modular Recombination. Science 301, 1904–1908 (2003).

2. Chen, Z. et al. Programmable design of orthogonal protein heterodimers. Nature 565, 106–111 (2019).

3. Chen, Z. et al. De novo design of protein logic gates. Science 368, 78–84 (2020).

4. Bragdon, M. D. J. et al. Cooperative assembly confers regulatory specificity and long-term genetic circuit stability. Cell 186, 3810–3825.e18 (2023).

5. Kalogriopoulos, N. A. et al. Synthetic GPCRs for programmable sensing and control of cell behaviour. Nature 637, 230–239 (2025).

6. Nelson, A. L., Dhimolea, E. & Reichert, J. M. Development trends for human monoclonal antibody therapeutics. Nat Rev Drug Discov 9, 767–774 (2010).

7. Lu, R.-M. et al. Development of therapeutic antibodies for the treatment of diseases. J Biomed Sci 27, 1 (2020).

8. Moutel, S. et al. NaLi-H1: A universal synthetic library of humanized nanobodies providing highly functional antibodies and intrabodies. eLife 5, e16228 (2016).

9. McMahon, C. et al. Yeast surface display platform for rapid discovery of conformationally selective nanobodies. Nat Struct Mol Biol 25, 289–296 (2018).

10. Wilson, P. C. & Andrews, S. F. Tools to therapeutically harness the human antibody response. Nat Rev Immunol 12, 709–719 (2012).

11. Cao, L. et al. Design of protein-binding proteins from the target structure alone. Nature 605, 551–560 (2022).

12. Bennett, N. R. et al. Atomically accurate de novo design of antibodies with RFdiffusion. Preprint at 10.1101/2024.03.14.585103 (2025).

13. Nabla Bio & Biswas, S. De novo design of epitope-specific antibodies against soluble and multipass membrane proteins with high specificity, developability, and function. Preprint at 10.1101/2025.01.21.633066 (2025).

14. Bennett, N. R. et al. Improving de novo protein binder design with deep learning. Nat Commun 14, 2625 (2023).

15. Baran, D. et al. Principles for computational design of binding antibodies. Proc. Natl. Acad. Sci. U.S.A. 114, 10900–10905 (2017).

16. Sormanni, P., Aprile, F. A. & Vendruscolo, M. Third generation antibody discovery methods: in silico rational design. Chem. Soc. Rev. 47, 9137–9157 (2018).

17. Muratspahić, E. et al. De novo design of miniprotein agonists and antagonists targeting G proteincoupled receptors. Preprint at 10.1101/2025.03.23.644666 (2025).

18. Whitehead, T. A. et al. Optimization of affinity, specificity and function of designed influenza inhibitors using deep sequencing. Nat Biotechnol 30, 543–548 (2012).

19. Dauparas, J. et al. Robust deep learning based protein sequence design using ProteinMPNN. (2023).

20. Jumper, J. et al. Highly accurate protein structure prediction with AlphaFold. Nature 596, 583–589 (2021).

21. Evans, R. et al. Protein complex prediction with AlphaFold-Multimer. Preprint at 10.1101/2021.10.04.463034 (2021).

22. Ravikumar, A., Arzumanyan, G. A., Obadi, M. K. A., Javanpour, A. A. & Liu, C. C. Scalable, Continuous Evolution of Genes at Mutation Rates above Genomic Error Thresholds. Cell 175, 1946–1957.e13 (2018).

23. Rix, G. et al. Continuous evolution of user-defined genes at 1 million times the genomic mutation rate. Science 386, eadm9073 (2024).

24. Wellner, A. et al. Rapid generation of potent antibodies by autonomous hypermutation in yeast. Nat Chem Biol 17, 1057–1064 (2021).

25. Paulk, A. M., Williams, R. L. & Liu, C. C. Rapidly Inducible Yeast Surface Display for Antibody Evolution with OrthoRep. ACS Synth. Biol. 13, 2629–2634 (2024).

26. Abramson, J. et al. Accurate structure prediction of biomolecular interactions with AlphaFold 3. Nature 630, 493–500 (2024).

27. Fleishman, S. J. et al. Computational Design of Proteins Targeting the Conserved Stem Region of Influenza Hemagglutinin. Science 332, 816–821 (2011).

28. Adams, R. M., Mora, T., Walczak, A. M. & Kinney, J. B. Measuring the sequence-affinity landscape of antibodies with massively parallel titration curves. eLife 5, e23156 (2016).

29. Starr, T. N. et al. Shifting mutational constraints in the SARS-CoV-2 receptor-binding domain during viral evolution. (2022).

30. Phillips, A. M. et al. Hierarchical sequence-affinity landscapes shape the evolution of breadth in an anti-influenza receptor binding site antibody. eLife 12, e83628 (2023).

31. Cao, L. et al. De novo design of picomolar SARS-CoV-2 miniprotein inhibitors. (2020).

32. Berger, S. et al. Preclinical proof of principle for orally delivered Th17 antagonist miniproteins. Cell 187, 4305–4317.e18 (2024).

33. Hie, B. L. et al. Efficient evolution of human antibodies from general protein language models. Nat Biotechnol 42, 275–283 (2024).

34. Shanker, V. R., Bruun, T. U. J., Hie, B. L. & Kim, P. S. Unsupervised evolution of protein and antibody complexes with a structure-informed language model. (2024).

35. Fournier, Q. et al. Protein Language Models: Is Scaling Necessary? Preprint at 10.1101/2024.09.23.614603 (2024).

36. Pisera, A. O., Yu, Y., Williams, R. L. & Liu, C. C. Ultra-efficient Integration of Gene Libraries onto Yeast Cytosolic Plasmids. ACS Synth. Biol. 14, 1002–1008 (2025).

37. Pisera, A. (Olek), Tanoori, A. & Liu, C. C. Rapid continuous evolution of gene libraries towards arbitrary functions. Preprint at 10.1101/2025.03.22.644768 (2025).

38. Van, M. V., Fujimori, T. & Bintu, L. Nanobody-mediated control of gene expression and epigenetic memory. Nat Commun 12, 537 (2021).

39. Wan, J. et al. High-throughput development and characterization of new functional nanobodies for gene regulation and epigenetic control in human cells. Preprint at 10.1101/2024.11.01.621523 (2024).

40. Sanford, A., Kiriakov, S. & Khalil, A. S. A Toolkit for Precise, Multigene Control in Saccharomyces cerevisiae. ACS Synth. Biol. 11, 3912–3920 (2022).

41. Gietz, R. D. & Schiestl, R. H. Frozen competent yeast cells that can be transformed with high efficiency using the LiAc/SS carrier DNA/PEG method. Nat Protoc 2, 1–4 (2007).

42. An, L. & Lee, G. R. De Novo Protein Design Using the Blueprint Builder in Rosetta. CP Protein Science 102, e116 (2020).

43. Leaver-Fay, A. et al. Rosetta3. in Methods in Enzymology vol. 487 545–574 (Elsevier, 2011).

44. McElroy, C. A., Dohm, J. A. & Walsh, S. T. R. Structural and Biophysical Studies of the Human IL-7/IL-7Rα Complex. Structure 17, 54–65 (2009).

45. Yao, S. et al. De novo design and directed folding of disulfide-bridged peptide heterodimers. Nat Commun 13, 1539 (2022).

46. Chao, G. et al. Isolating and engineering human antibodies using yeast surface display. Nat Protoc 1, 755–768 (2006).

47. Benatuil, L., Perez, J. M., Belk, J. & Hsieh, C.-M. An improved yeast transformation method for the generation of very large human antibody libraries. Protein Engineering, Design and Selection 23, 155–159 (2010).

48. Estes, B. et al. Next generation Fc scaffold for multispecific antibodies. iScience 24, 103447 (2021).

49. Buschmann, T. & Bystrykh, L. V. Levenshtein error-correcting barcodes for multiplexed DNA sequencing. BMC Bioinformatics 14, 272 (2013).

50. Chen, S., Zhou, Y., Chen, Y. & Gu, J. fastp: an ultra-fast all-in-one FASTQ preprocessor. Bioinformatics 34, i884–i890 (2018).

51. Zhang, J., Kobert, K., Flouri, T. & Stamatakis, A. PEAR: a fast and accurate Illumina Paired-End reAd mergeR. Bioinformatics 30, 614–620 (2014).

52. Langmead, B. & Salzberg, S. L. Fast gapped-read alignment with Bowtie 2. Nature Methods 9, 357–359 (2012).

53. Virtanen, P. et al. SciPy 1.0: fundamental algorithms for scientific computing in Python. Nat Methods 17, 261–272 (2020).

54. Hagberg, A. A., Schult, D. A. & Swart, P. J. Exploring Network Structure, Dynamics, and Function using NetworkX. in 11–15 (Pasadena, California, 2008). doi:10.25080/TCWV9851.

55. Pedregosa, F. et al. Scikit-learn: Machine Learning in Python. MACHINE LEARNING IN PYTHON.

56. Phillips, A. M. et al. Binding affinity landscapes constrain the evolution of broadly neutralizing antiinfluenza antibodies. eLife 10, e71393 (2021).

57. Heyne, M., Papo, N. & Shifman, J. M. Generating quantitative binding landscapes through fractional binding selections combined with deep sequencing and data normalization. Nat Commun 11, (2020).

58. Tinberg, C. E. et al. Computational design of ligand-binding proteins with high affinity and selectivity. Nature 501, 212–216 (2013).

59. Gaspar, J. M. NGmerge: merging paired-end reads via novel empirically-derived models of sequencing errors. BMC Bioinformatics 19, (2018).

60. Huang, H., Wang, Y. & Rudin, C. Navigating the Effect of Parametrization for Dimensionality Reduction.

61. Tareen, A. & Kinney, J. B. Logomaker: beautiful sequence logos in Python. Bioinformatics 36, 2272–2274 (2020).

