## Supplementary material for "Mapping the evolution of computationally designed protein binders": Figure S

### Supplementary Text

#### A pre-trained protein language model can be used to generate additional candidate sequences for Mb-4

We explored the feasibility of integrating machine learning with our evolutionary campaigns to guide future diversification efforts. Specifically, we used a pre-trained 350M-parameter protein language model (AMPLIFY-350M) to perform in silico sampling around the parent sequence of the Mb-4 neutral drift outcome library (Figure S18A). These simulations were designed to approximate neutral drift by conditioning amino acid probabilities on local sequence context, and they revealed patterns that both mirrored and differed from observed mutational outcomes in the OrthoRep-derived library. For example, while AMPLIFY-350M generally predicted uniform substitution preferences at highly mutated positions, certain residues retained strong wild-type bias (e.g., 31E compared to 40G; Figure S18B), suggesting model-informed constraints consistent with evolutionary conservation. To assess the utility of this modeling approach for predicting functional outcomes, we trained a simple classifier on AMPLIFY-derived sequence embeddings and binary binding labels from the OrthoRep dataset. The model achieved modest but meaningful predictive performance across leave-one-residue-out validation splits (Fig. S19C and S19D). Notably, projecting both experimentally observed and in silico-generated variants into a two-dimensional fitness landscape revealed that the computationally sampled sequences populate unexplored regions of sequence space that flank and extend beyond the OrthoRep library (Figure S19A). These early results suggest that protein language models, when paired with simple downstream predictors trained on experimental data such as those generated by OrthoRep-driven evolution, could help prioritize starting points for future evolution campaigns and support active learning frameworks for protein binder discovery.

### Supplementary Methods

#### Sampling sequences from the pre-trained AMPLIFY protein language model

AMPLIFY-350M (weights downloaded from Hugging Face) was used to sample candidates around the parent sequence in a protocol closely mimicking neutral drift. The inputs to the sampling algorithm were a parent sequence (of length say L), a list of residue positions (say P) to mutate and the desired number of samples (say N). |P| denotes the number of positions in P. To sample an amino acid $j$ $(1 \le j \le 20)$, at residue position $i$ in the sequence, first the amino acid at $i$ is masked out and the resulting sequence is passed through AMLIFY-350M to obtain a L×20 matrix $A$, where the $A_{ij}$ represents the probability of amino acid $j$ at residue position $i$. Then the amino acid is replaced with another one drawn from the distribution of all amino acids at that position— i.e., from the $i_{th}$ row of matrix $A$. The sampling process is repeated for |P| successive iterations, where the position $i$ from P was selected at random in each iteration, and the parent sequence was replaced by the mutated sequence at the end of that iteration. Here, each iteration can be thought of as a computational "generation" and a sample is recorded as the mutated sequence at the end of |P| generations. Figure S18 shows a parallelized scheme used to generate N such samples, efficiently. Language model embeddings of sampled sequences used to generate Figure S19A were extracted from the last attention layer of AMPLIFY-350M and mean pooled across the sequence length to produce consistently shaped (960-dimensional vector) sequence representations. The sampling algorithm was implemented in Python v3.10.18 and AMPLIFY model inference was carried out on a single L40S GPU core.

The 2-D fitness landscape in Figure S19A was produced through a PaCMAP transformation of the higher dimensional sequence embeddings with parameters "n_components = 2, n_neighbors = 100, MN_ratio = 0.5, FP_ratio = 2.0, random_state = 42". For a fixed random_state, we observed no major change in the output landscape when n_neighbors was changed to 50, or when MN_ratio and FP_ratio were each varied between 0.5 and 3.

To predict the effects of novel mutations on Mb-4's binding affinity, we trained machine learning models using sequence embeddings generated by AMPLIFY-350M. Full-length minibinder sequences were passed through the model, and mean pooling was applied to the resulting embeddings to produce fixed-length feature vectors.

These pooled embeddings were used as input to a logistic regression classifier implemented with scikit-learn v1.7.0, using L2-regularization (regularization strength = 1). The classifier outputs a binary label indicating whether a mutant exhibits improved binding relative to the parent sequence, as determined experimentally (Figure 3).

Model training was conducted on neutral drift data using a leave-one-residue-out cross-validation scheme where residues with 10 to 100 mutants were used for validation. For each of 28 individual residue positions that met these sample size criteria, mutants with substitutions at that position were excluded from training and reserved for testing. This strategy enabled evaluation of the model's ability to generalize to previously unseen mutation sites. A final model was subsequently trained on the complete neutral drift dataset.

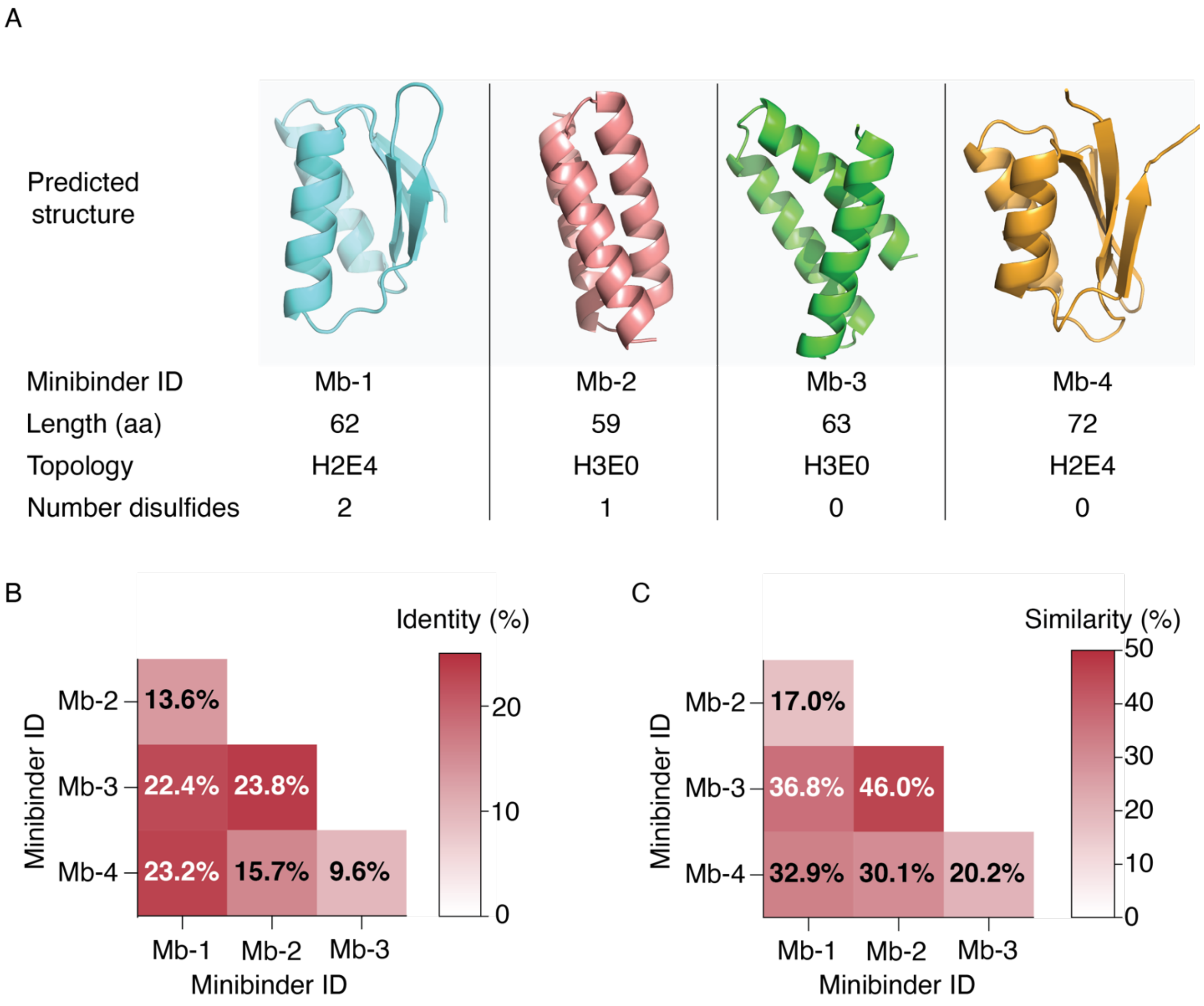

**Figure S1. The computationally designed minibinders are diverse in their sequences and design topology. A,** AlphaFold3 predicted structures of the computationally-designed minibinders and their associated features. **B and C,** Pairwise identity and similarity heatmaps. Alignments were computed using the Needleman-Wunsch algorithm and the metrics were calculated based on the BLOSUM62 matrix.

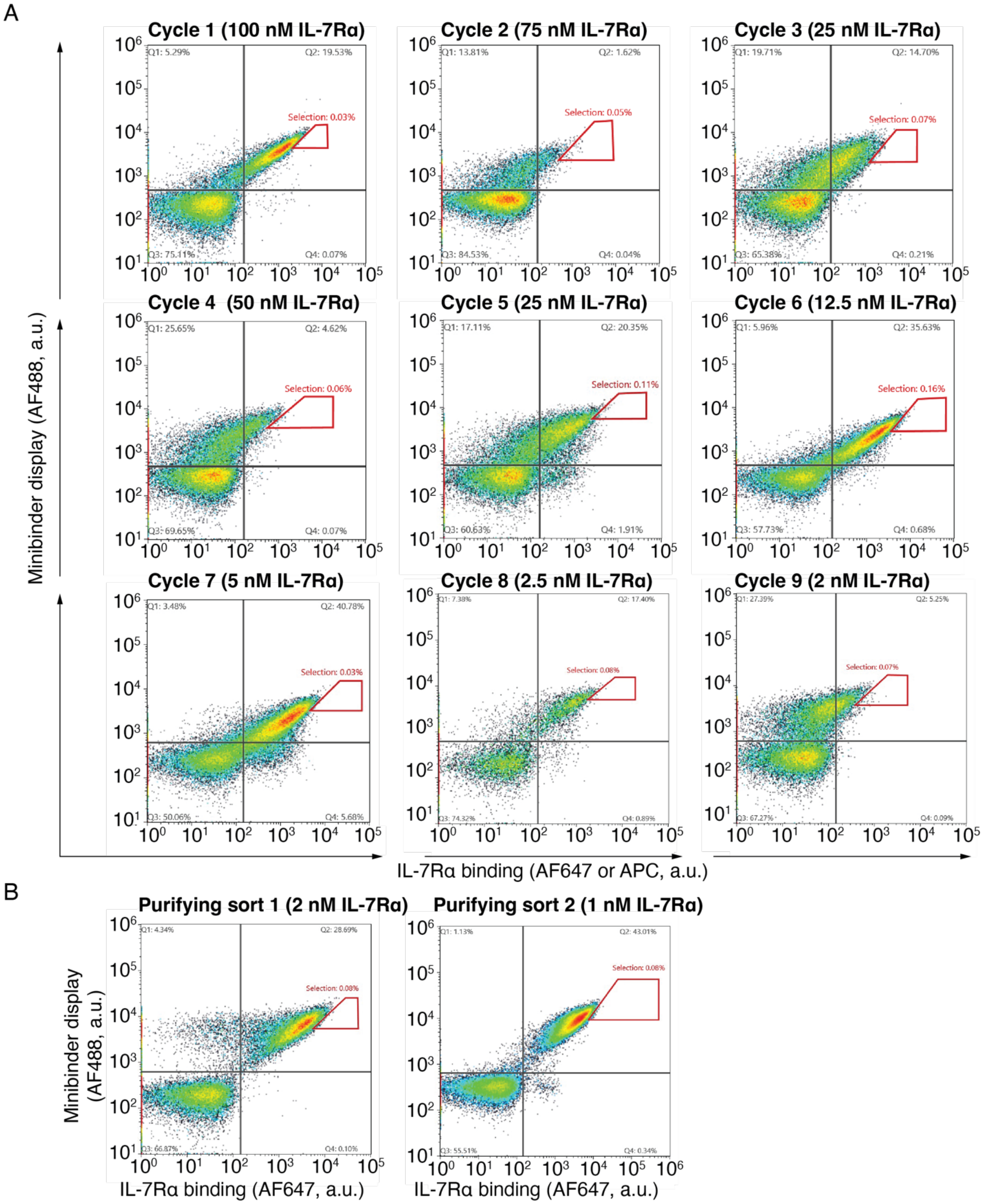

**Figure S2. Mb-1 was affinity matured using AHEAD. A,** Sequential FACS plots showing affinity maturation of Mb-1. For cycles 1 and 3, we used streptavidin-APC with a biotinylated human IL-7Rα. In all other cycles, we used anti-human IgG-AF647 with IgG-Fc tagged human IL-7Rα. **B,** Purifying sorts used to isolate clones expressing the best binders. The red polygons represent the gates used for selecting the best binders.

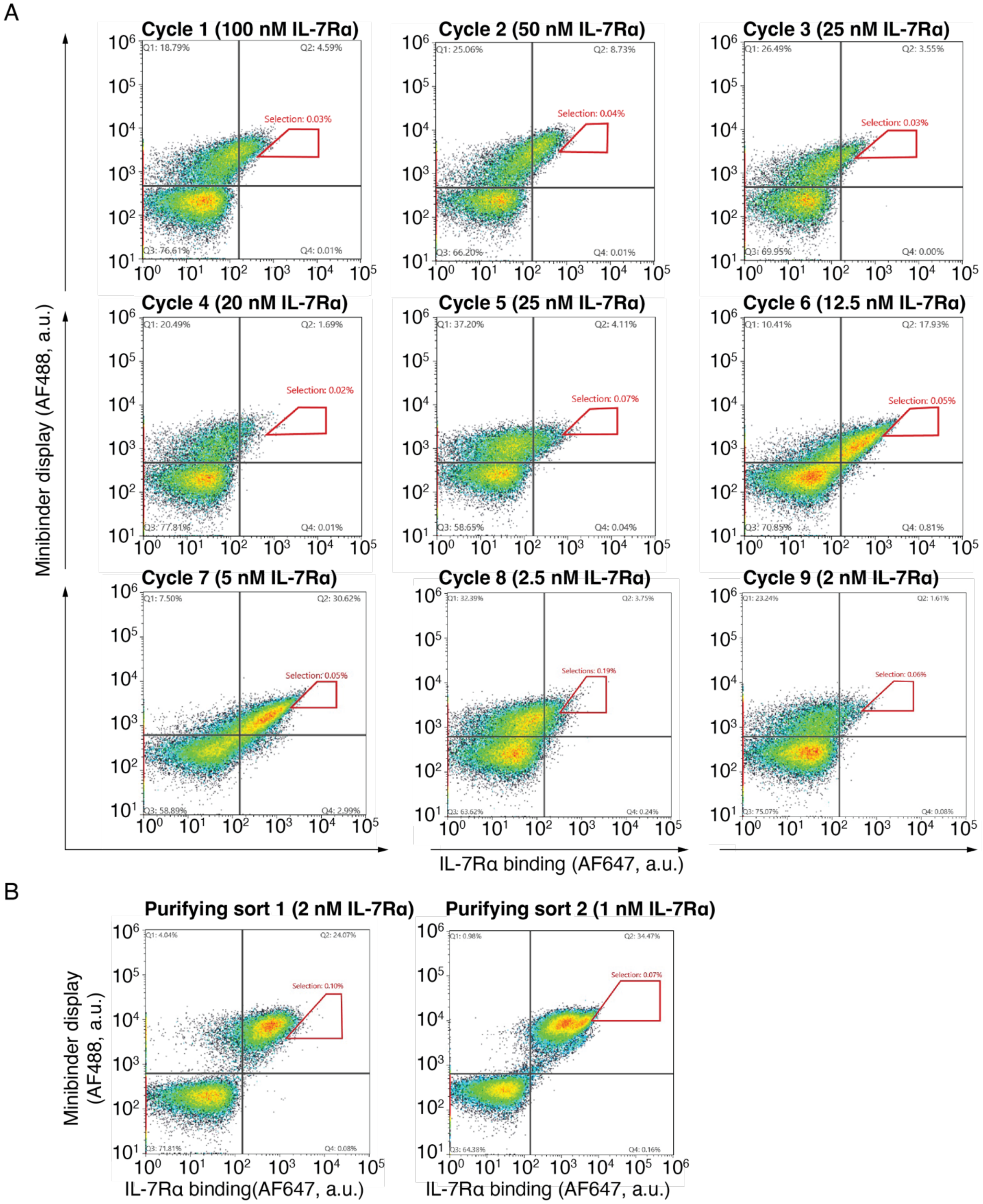

**Figure S3. Mb-2 was affinity matured using AHEAD. A,** Sequential FACS plots showing affinity maturation of Mb-2. **B,** Purifying sorts used to isolate clones expressing the best binders. The red polygons represent the gates used for selecting the best binders.

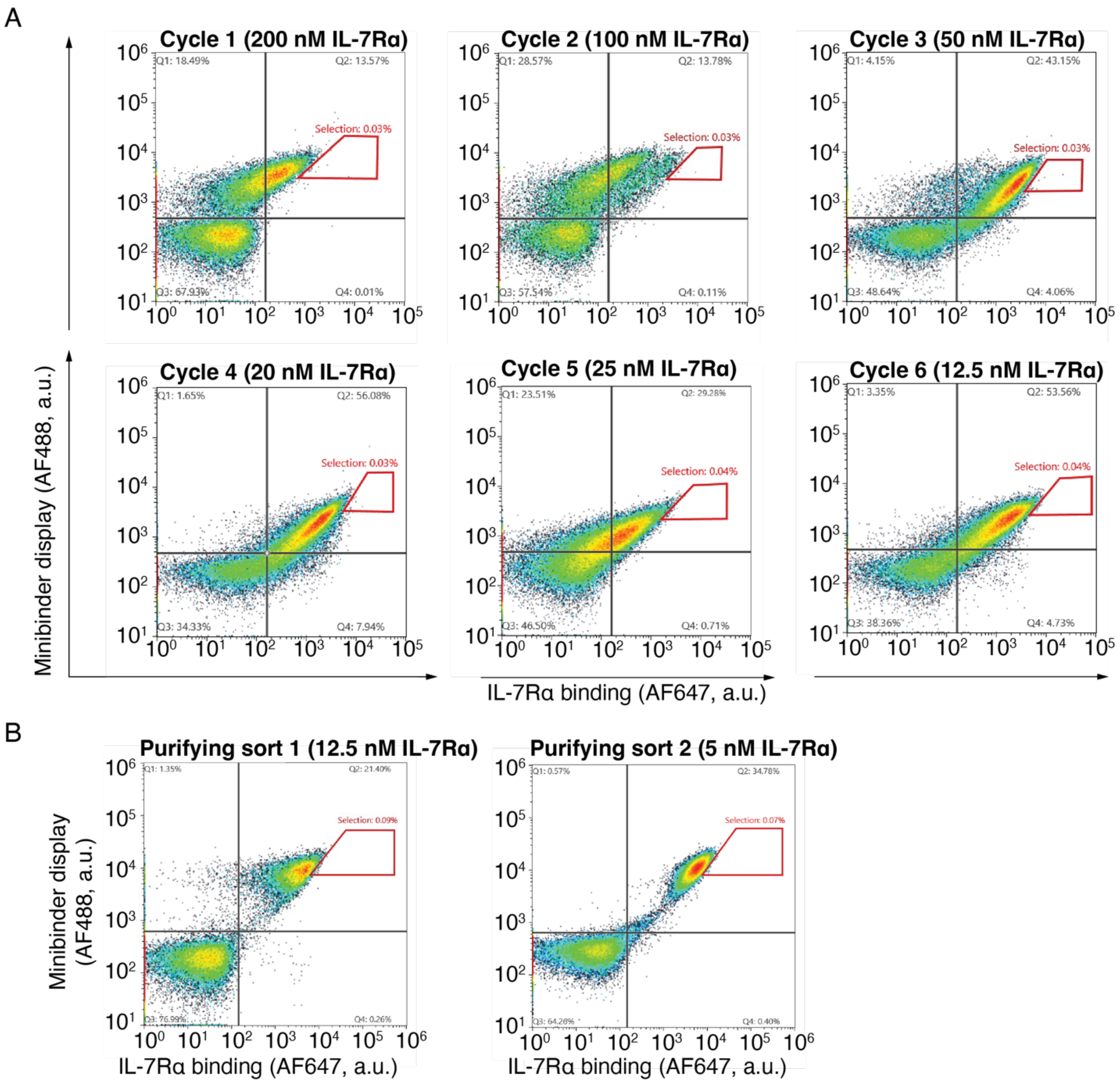

**Figure S4. Mb-3 was affinity matured using AHEAD. A,** Sequential FACS plots showing affinity maturation of Mb-3. **B,** Purifying sorts used to isolate clones expressing the best binders. The red polygons represent the gates used for selecting the best binders.

A

B

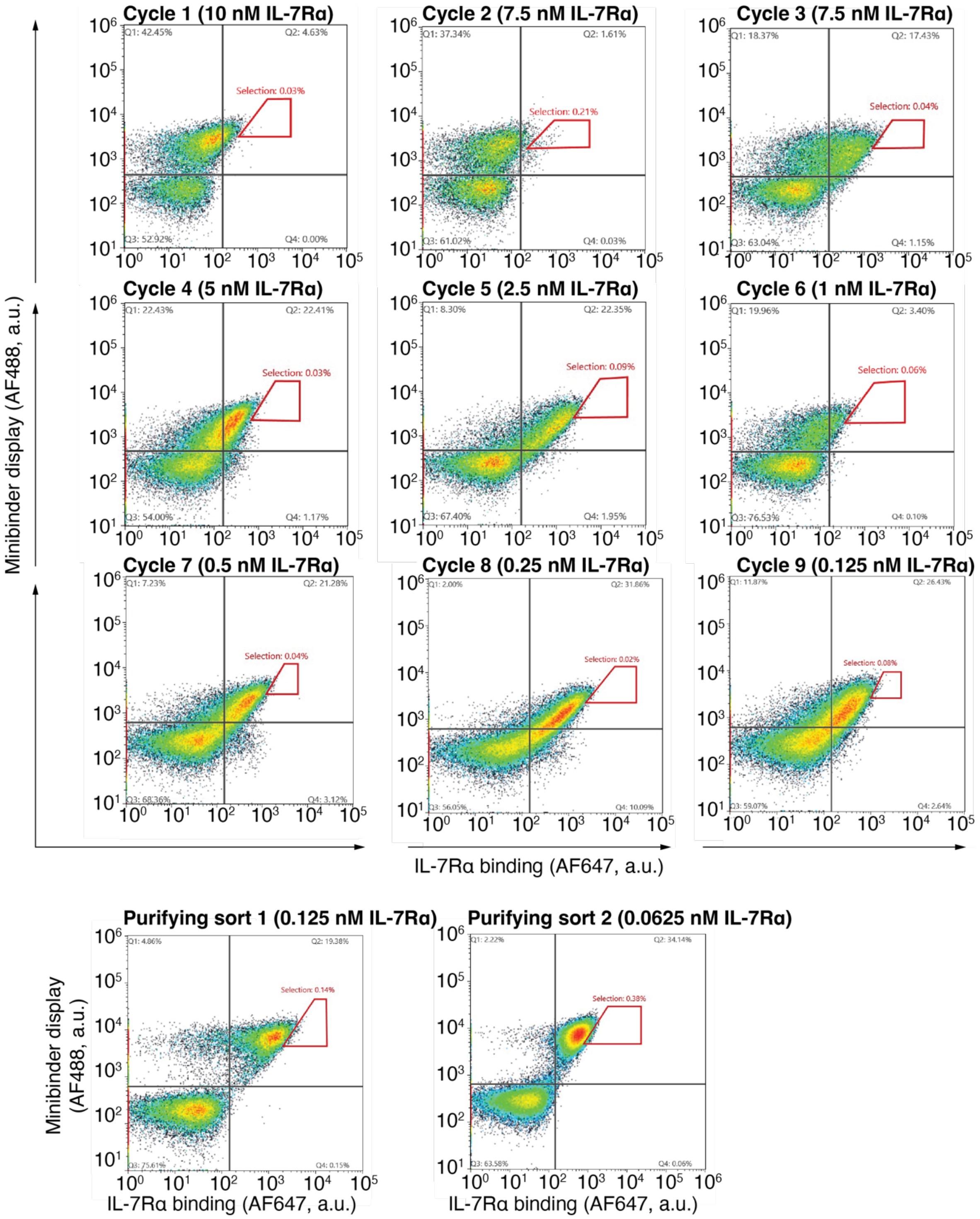

**Figure S5. Mb-4 was affinity matured using AHEAD. A,** Sequential FACS plots showing affinity maturation of Mb-4. **B,** Purifying sorts used to isolate clones expressing the best binders. The red polygons represent the gates used for selecting the best binders.

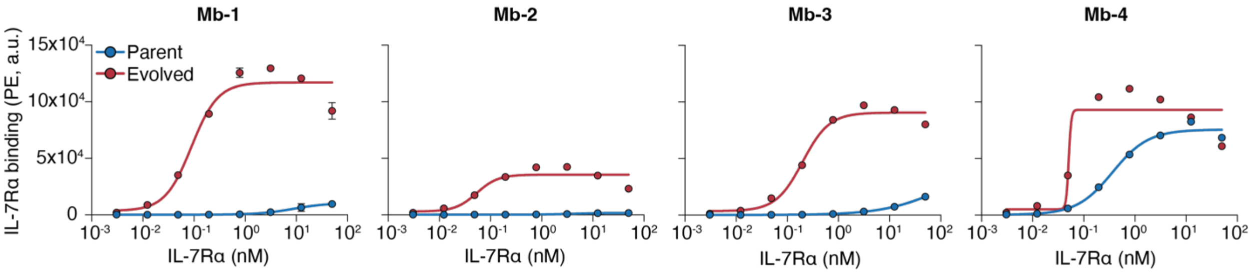

**Figure S6. AHEAD improves binding for Mb-1 through Mb-4.** Shown are dose-response curves for yeast displayed parent (blue) and evolved (red) minibinders. IL-7Rα binding for each biological replicate is taken as the median PE intensity for the displaying population. Each point represents median IL-7Rα binding ± one standard deviation for n=2-3 biological replicates.

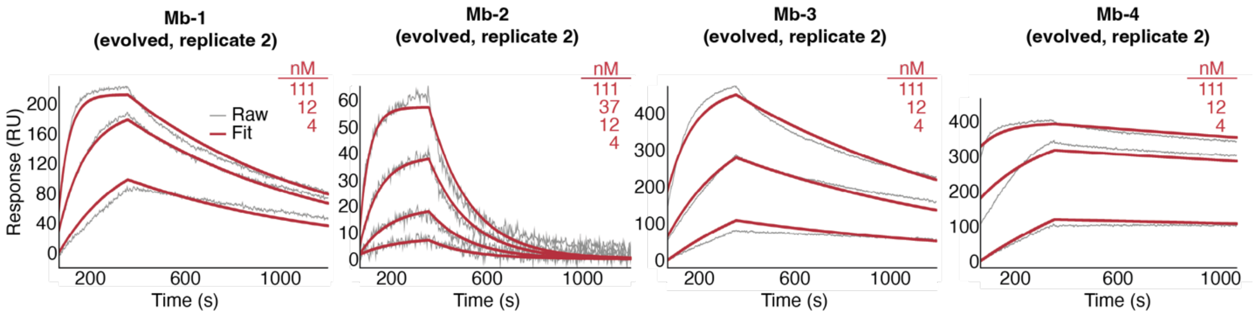

**Figure S7. Additional SPR traces for the evolved minibinders.** SPR traces for the second biological replicate for the affinity matured minibinders.

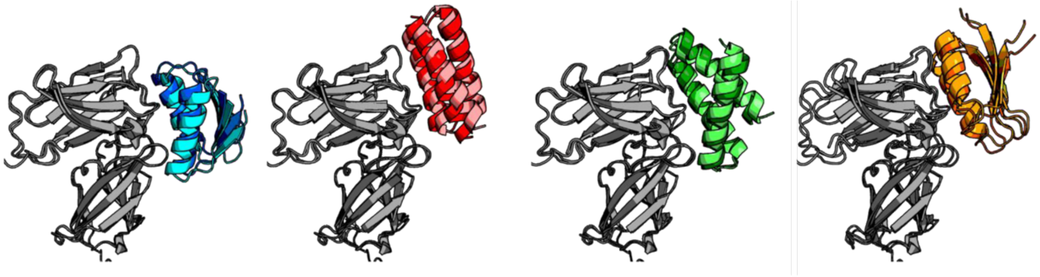

**Figure S8. Structure predictions produce a consistent target binding mode.** Alignment of AF2-Multimer and AF3 predicted complex structures. Models were aligned on IL-7Rα Cα atoms (grey) and Cα RMSD of minibinders were calculated with no fitting, in the range of 2.2-2.9 Å. Minibinder-only heavy atom RMSD with fitting was 1.2-1.3 Å. AF3 models in lighter colors, AF2 models in darker colors; from left to right: Mb-1, Mb-2, Mb-3, Mb-4.

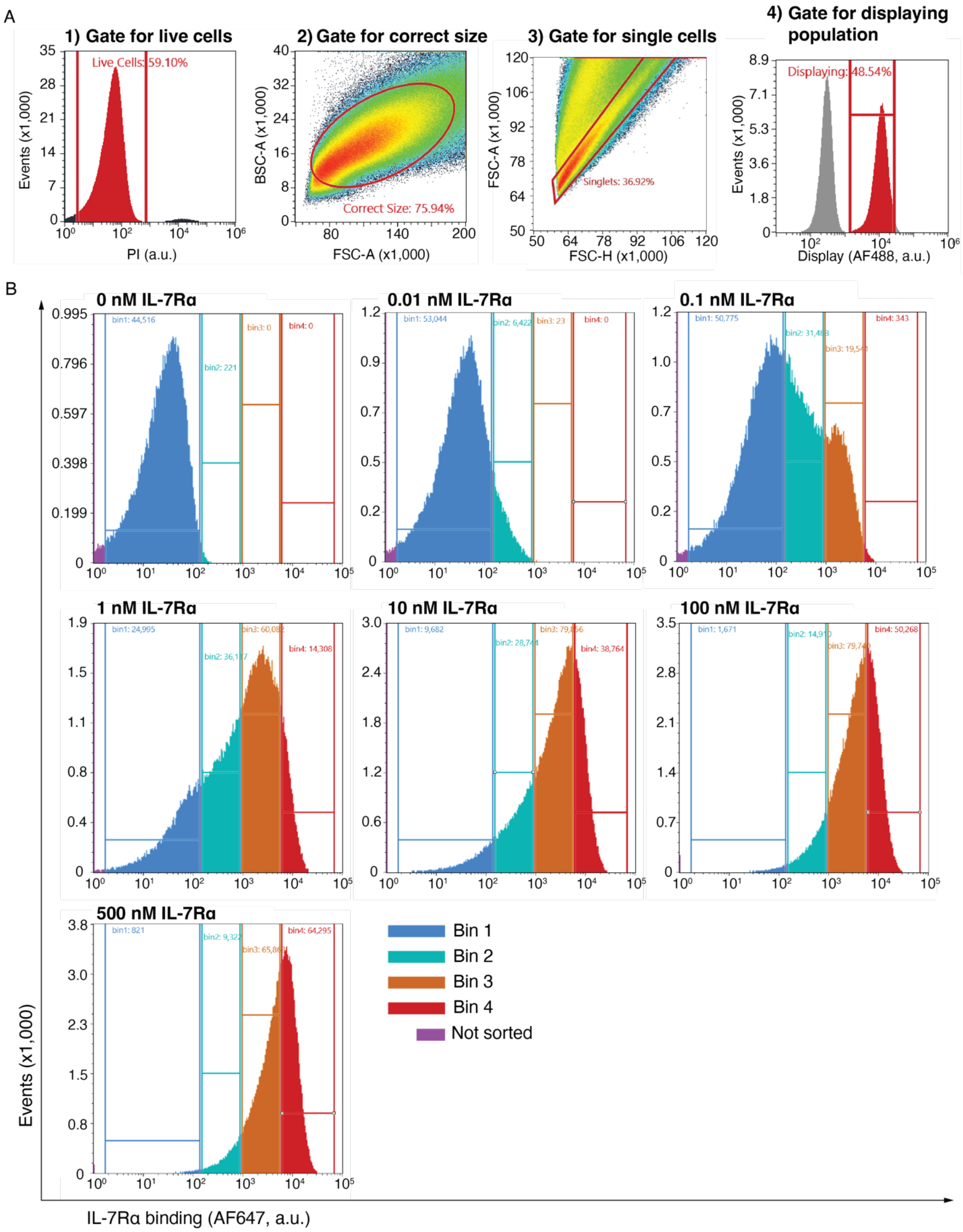

**Figure S9. The gating strategy for FACS selections in the Tite-seq assay. A,** Representative gating strategy for the FACS selections. First, live cells were selected based on the presence of propidium iodide (PI) signal. Next, yeast cells were selected based on FSC-A and BSC-A. Single yeast cells were then isolated using FSC-H and FSC-A. Lastly, were focused our analysis on yeast cells that were displaying minibinder. **B,** Raw binding distributions and gating thresholds at the seven antigen concentrations used during the Tite-seq assay for one representative biological replicate.

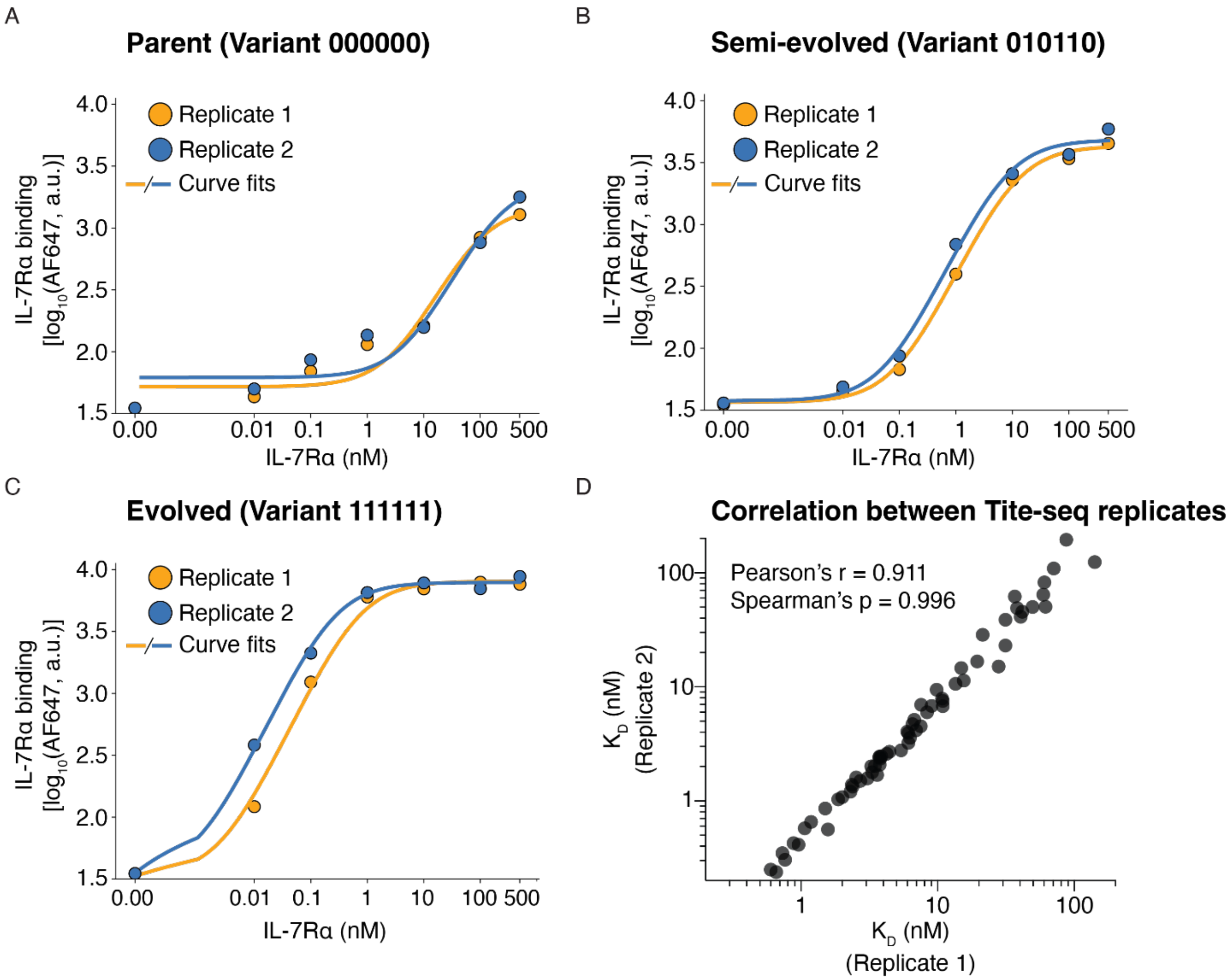

**Figure S10. Tite-seq can be used to characterize the combinatorial landscape of Mb-1. A-C,** Dose-response curves extracted from the Tite-seq assay for the parent, semi-evolved, and evolved minibinder variants. The minibinder identifiers (e.g., 010110) correspond to those shown in Figure 2. The x-axis is plotted on a log-scale, except for the domain between 0-0.001 nM which is plotted on a linear scale. **D,** Monovalent dissociation values ($K_D$) obtained from two biological replicates.

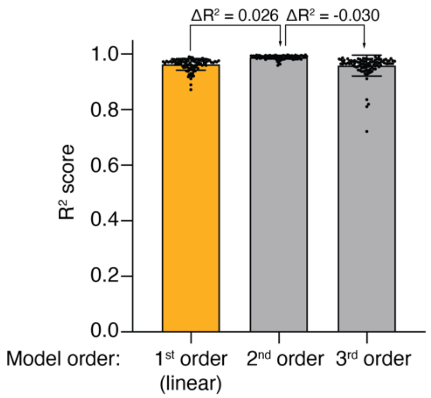

**Figure S11. A linear model is sufficient to explain mutational effects in Mb-1.** Shown are the $R^2$ scores for test set predictions in 100 different train-test splits.

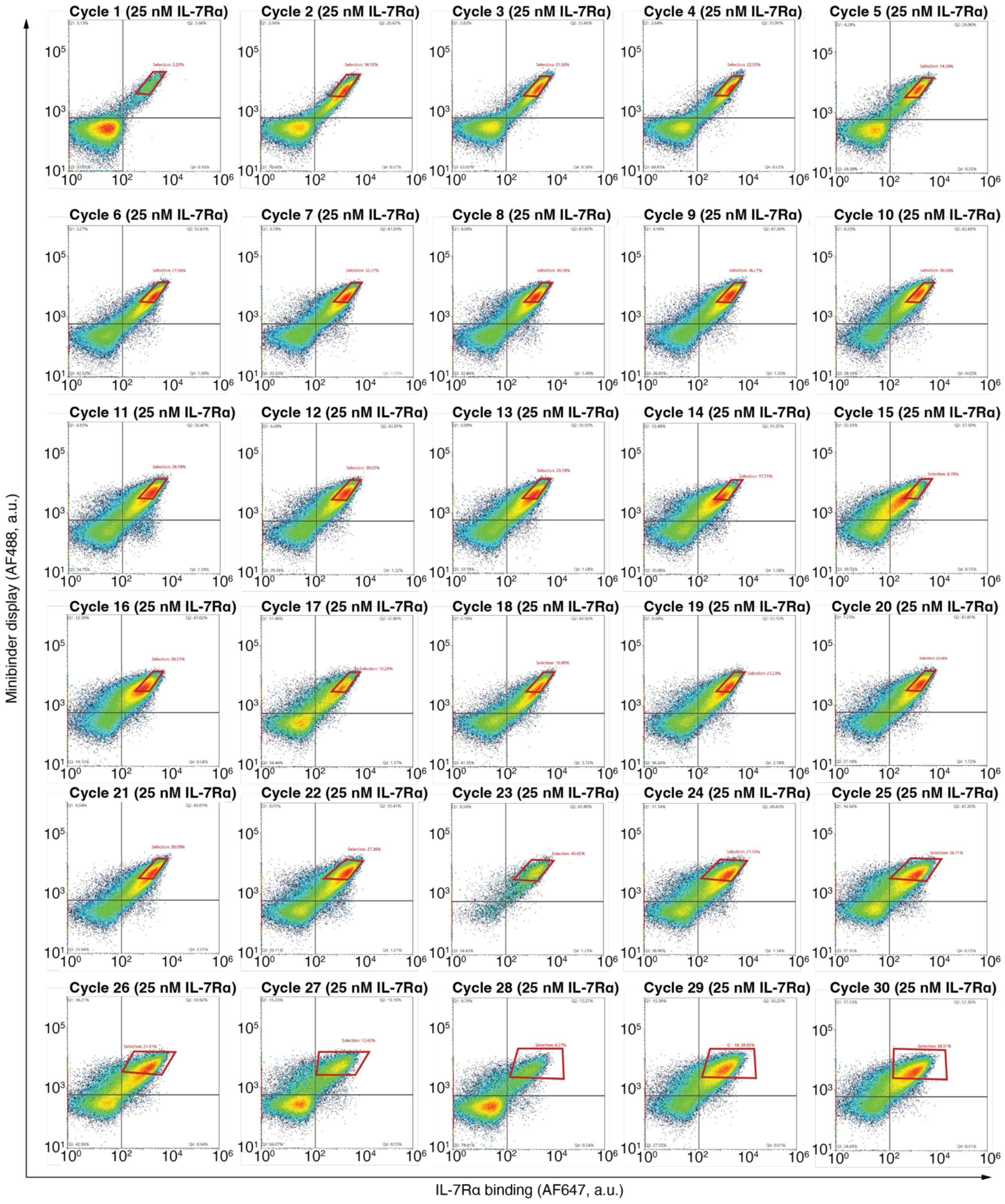

**Figure S12. Mb-4 was diversified using modified AHEAD cycles.** Sequential FACS plots showing the diversification of Mb-4. The red polygons represent the gates used for selecting functional binders. The gates were redrawn in Illustrator (Adobe) to make them more visible.

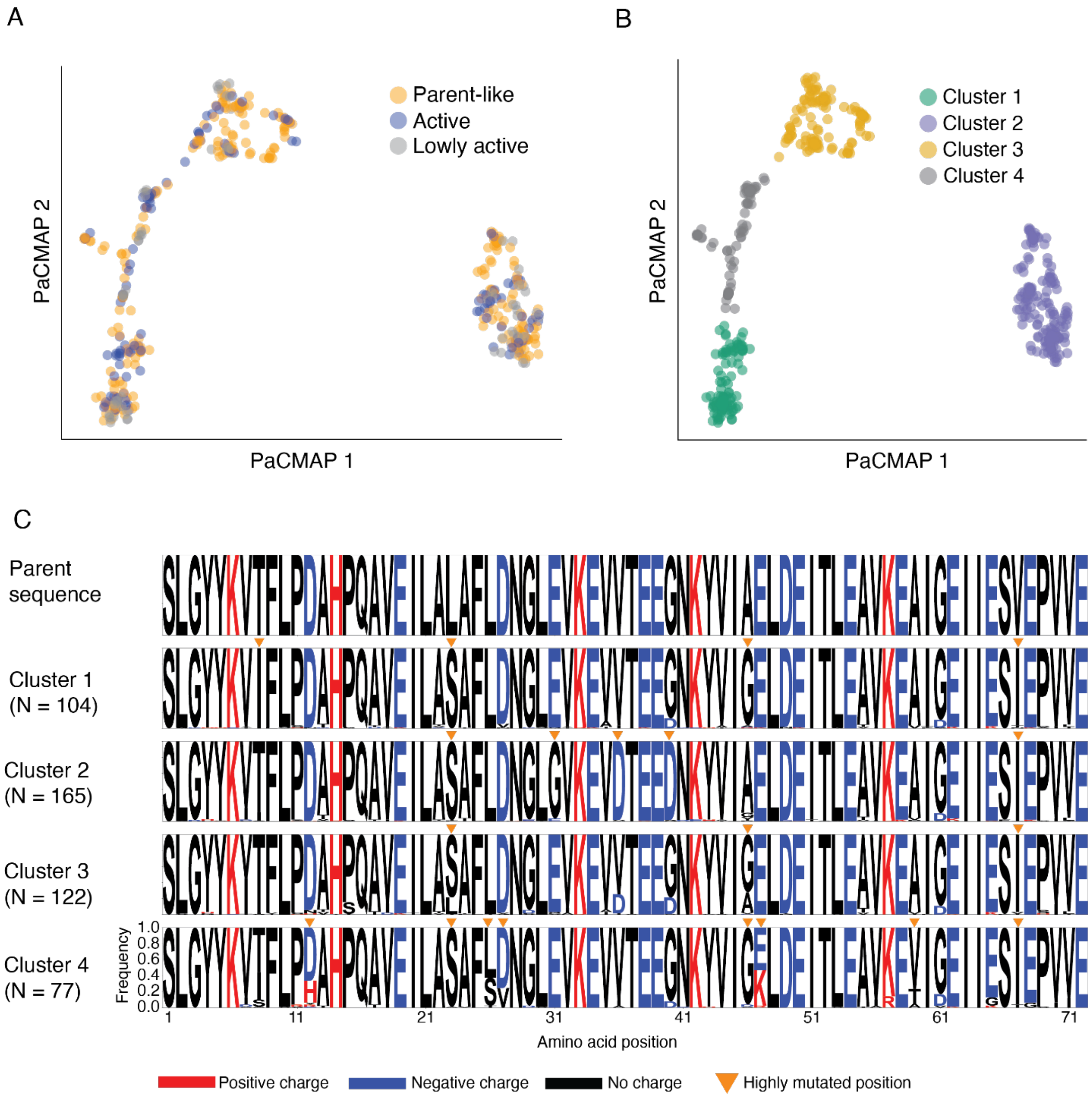

**Figure S13. Diversified Mb-4 library contains distinct sequence clusters. A,** Two-dimensional representations for all unique protein sequences obtained using PACMap. Sequences are colored by the activity class labels shown in Figure 3. **B,** The PACMap shown in panel A, with sequences clustered using a K-means clustering algorithm with K = 4. **C,** Sequence logos for the clusters obtained in panel B and the parent sequence for reference. The orange triangles indicate positions where a high proportion of sequences in a cluster contain a mutation, relative to the parent sequence.

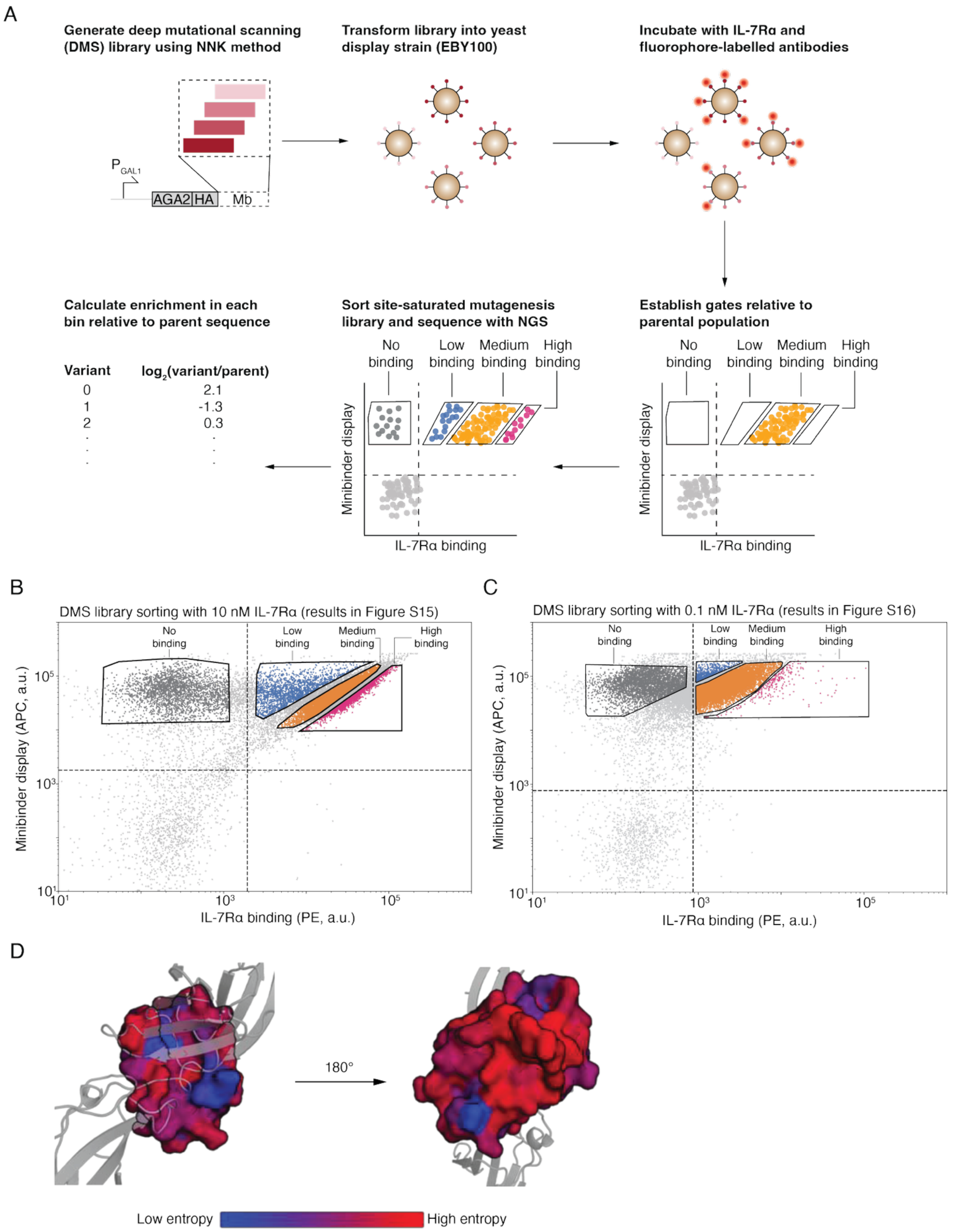

**Figure S14. A deep mutational scanning (DMS) library was used to characterize the impact of single amino acid substitutions on binding for Mb-4. A,** Schematized is the workflow used to generate and analyze the Mb-4 DMS library. **B and C,** FACS plot showing the distribution of binding for Mb-4 DMS library incubated with **(B)** 10 nM and **(C)** 0.1 nM IL-7Rα, and the thresholds used to sort cells based on binding. **D,** Positional Shannon entropy of the DMS enrichment normalized to wild type for Mb-4 was calculated and mapped to AF2 model for visualization. Low entropy (bluer) indicates high conservation of residues predicted to contact IL-7Rα. Higher entropy (redder) was observed in regions predicted distant from the receptor and not in the minibinder hydrophobic core.

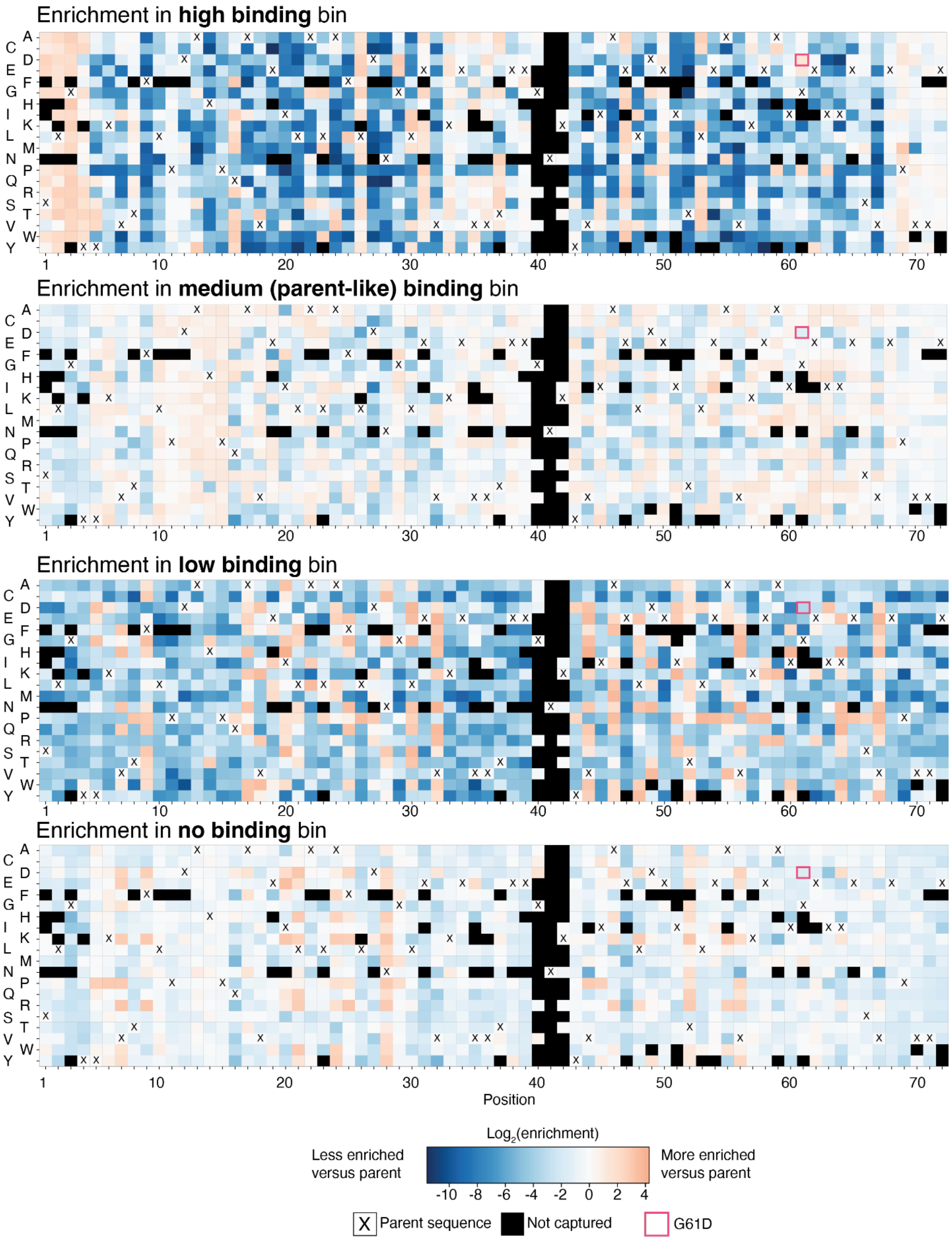
Enrichment in **high binding** bin
Enrichment in **medium (parent-like) binding** bin
Enrichment in **low binding** bin
Enrichment in **no binding** bin
A C D E F G H I K L M N P Q R S T V W Y
1 10 20 30 40 50 60 70
Position
Log2(enrichment)
Less enriched versus parent
More enriched versus parent
-10 -8 -6 -4 -2 0 2 4
X Parent sequence
Not captured
G61D

**Figure S15. Deep mutational scanning elucidates the impact of single amino acid mutations on binding for Mb-4 with 10 nM IL-7Rα.** Heatmaps show enrichment of minibinders with single mutations versus the parent sequence in high binding, medium (parent-like) binding, low binding, and no binding bins. Yeast displaying the minibinder library were incubated with 10 nM IL-7Rα.

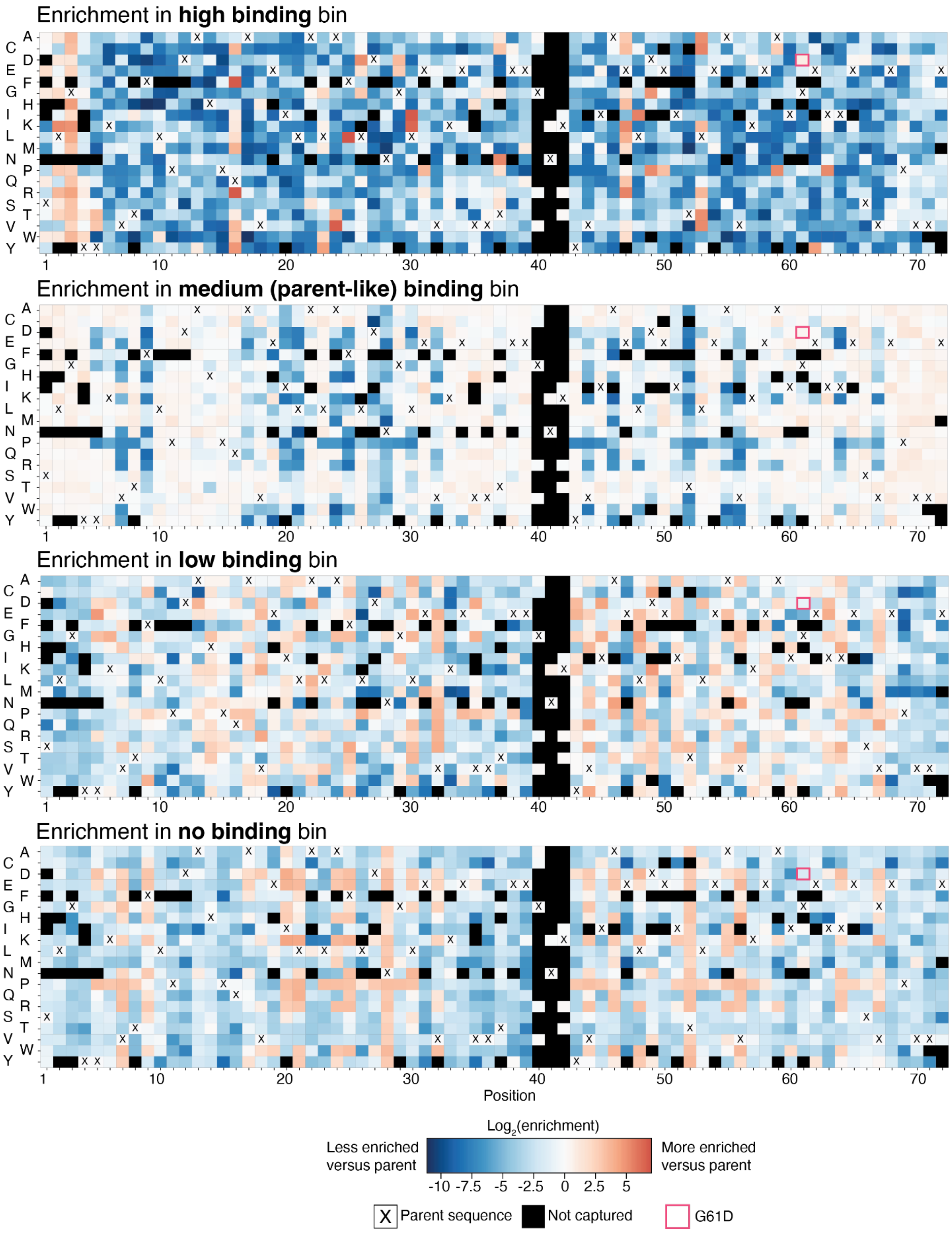

**Figure S16. Deep mutational scanning elucidates the impact of single amino acid mutations on binding for Mb-4 with 0.1 nM IL-7Rα.** Heatmaps show enrichment of minibinders with single mutations versus the parent sequence in high binding, medium (parent-like) binding, low binding, and no binding bins. Yeast displaying the minibinder library were incubated with 0.1 nM IL-7Rα.

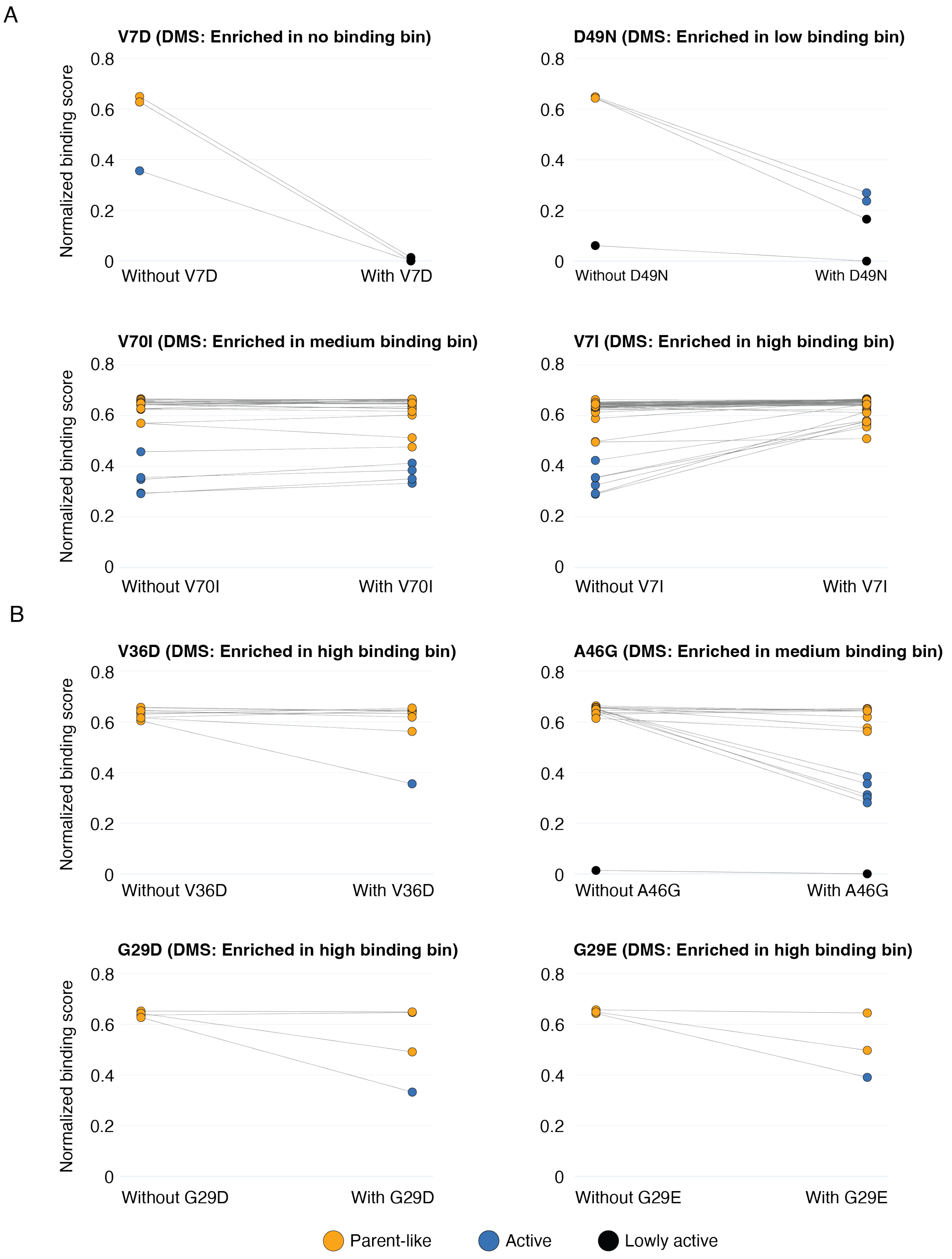

**Figure S17. Comparison between matched Mb-4 sequences differing by a single amino acid mutation. A,** Mutations that agree with DMS results. **B,** Mutations that appear to have context-dependent effects.

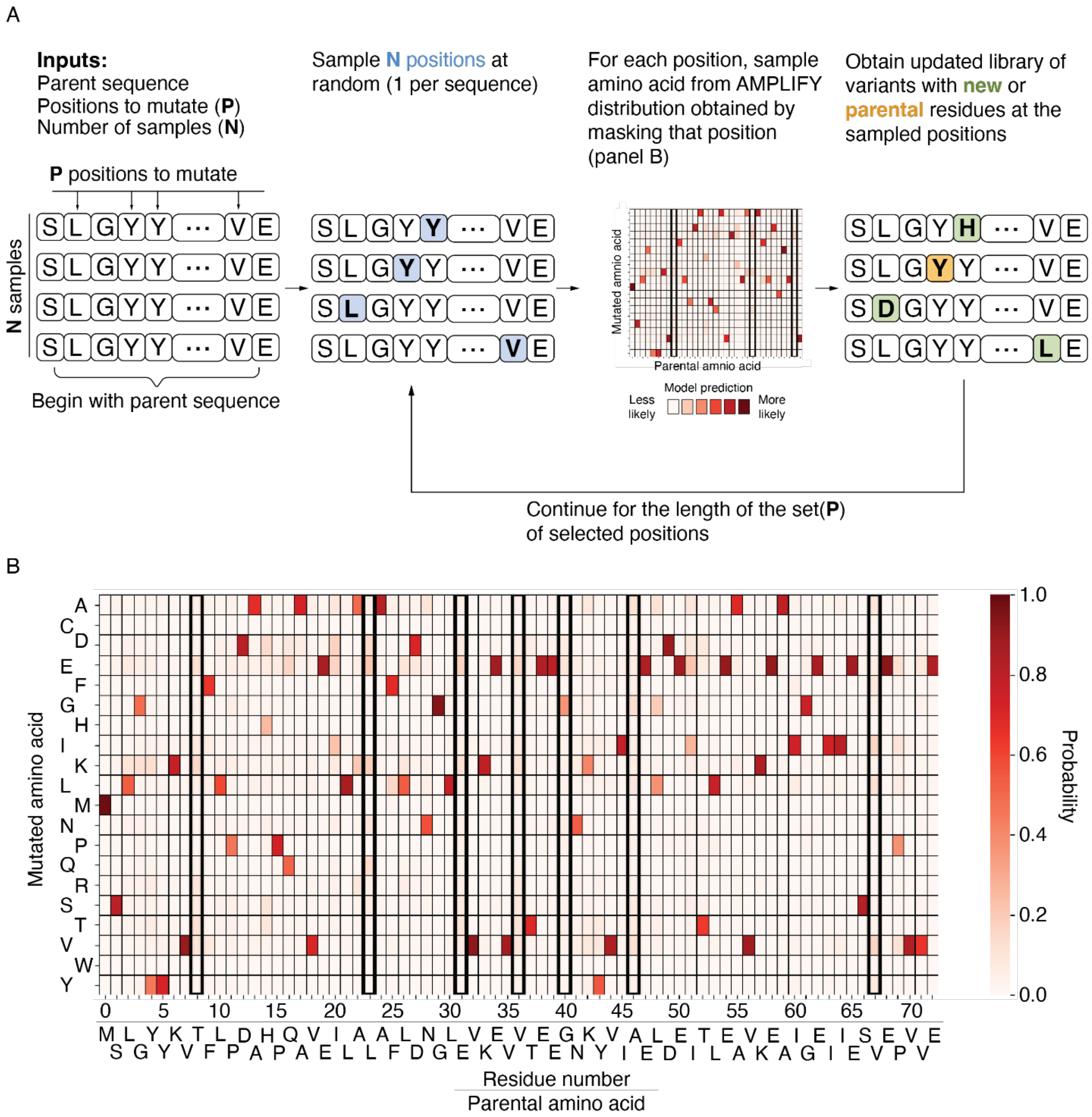

**Figure S18. Parallelized sampling algorithm for generating sequences around a given parent from the pre-trained protein language model AMPLIFY.** For a given parent sequence, a set of positions to mutate **P,** and the required number of samples (N), each iteration involves picking N residue positions at random, masking the amino acids at these positions, passing the masked sequences through AMPLIFY-350M, and replacing the amino acids at these positions by sampling randomly from the probability matrices produced from AMPLIFY-350M. Continuing this for |P| iterations (|P| = length of the set **P**), produces N samples of varying diversity from the starting parent. **B,** Amino acid substitution probabilities determined by AMPLIFY-350M. The boxed columns correspond to residues with uniform substitution preferences.

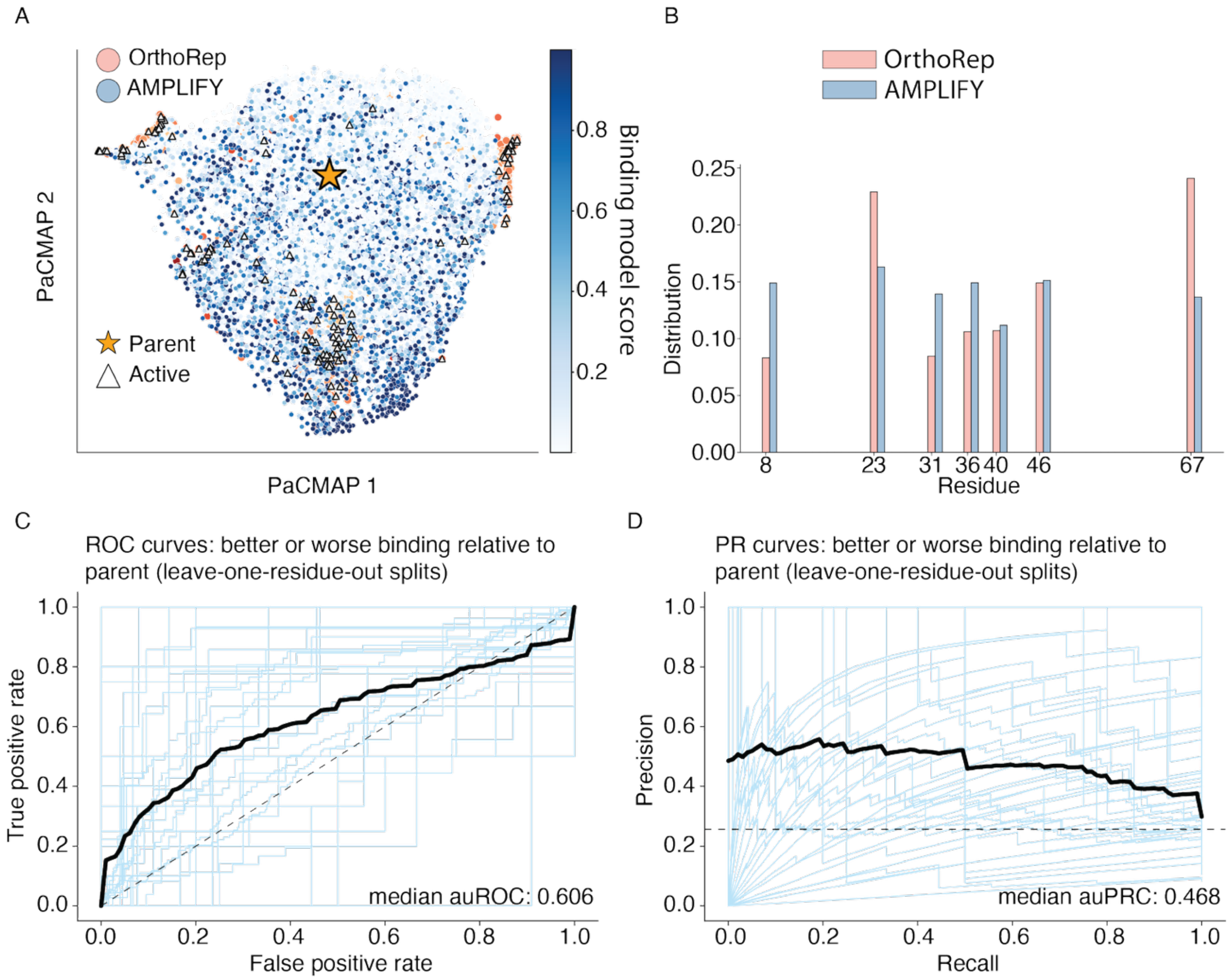

**Figure S19. Sequence space exploration by machine-learning models trained from the diversified minibinder library. A.** 2-D Fitness landscape of AMPLIFY sample embeddings, colored according to the score attributed to them from a simple binding model trained on the experimental library. In-silico predicted active candidates (dark blue) from AMPLIFY samples have some overlap with active candidates from the OrthoRep library (marked by triangles, towards the middle) but can be distant in sequence space from the parent and from other library members as well (lower right boundary). **B.** Distribution of residues mutated in 8000 sample sequences generated from AMPLIFY (see Supplemental Methods) around the Mb-4 parent. **C,** Receiver operating characteristic (ROC) and **D,** precision-recall (PR) curves for leave-one-residue-out cross-validation splits for binding model evaluation. Results from individual splits are shown in blue and the median curves are shown in black. For the ROC curve (panel C), the dashed line corresponds to the performance of a random classifier. For the PR curve (panel D), the dashed line corresponds to the prevalence of variants with better binding compared to the parent.

**Table S1. Sequences for the minibinders and IL-7Rα used in this study.** For human IL-7Rα, bold indicates the extracellular domain that was recombinantly expressed, purified, and used in this study.

| **Sequence ID** | **Amino acid sequence** |
|---|---|
| Mb-1 (parent) | CCLTFENPEFAEQAVLIAEDAGLKVEVKPGEDSLTVCLSNEAACKYFAERA<br>KEMGVPVTVEP |
| Mb-2 (parent) | SAEDELWELLDECRAAYFEGKFKEAKKCLKKVLELAKKNGNKAFEEAAK<br>ALLKAVEAAE |
| Mb-3 (parent) | SMFEEMVEKMIEYLKACKESDYDKADDLYGEIIDLADEAAKGNSEKAREL<br>IEKAYEEAKKRYE |
| Mb-4 (parent) | SLGYYKVTFLPDAHPQAVEILALAFLDNGLEVKEVVTEEGNKYVIAELDEI<br>TLEAVKEAIGEIIESVEPVVE |
| Human IL-7Rα (GenBank accession AAC83204) | MTILGTTFGMVFSLLQVVSG**ESGYAQNGDLEDAELDDYSFSCYSQLEVN**<br>**GSQHSLTCAFEDPDVNTTNLEFEICGALVEVKCLNFRKLQEIYFIETKK**<br>**FLLIGKSNICVKVGEKSLTCKKIDLTTIVKPEAPFDLSVIYREGANDFVV**<br>**TFNTSHLQKKYVKVLMHDVAYRQEKDENKWTHVNLSSTKLTLLQRK**<br>**LQPAAMYEIKVRSIPDHYFKGFWSEWSPSYYFRTPEINNSSGEMD**PILLI<br>ISILSFFSVALLVILACVLWKKRIKPIVWPSLPDHKKTLEHLCKKPRKNLNV<br>SFNPESFLDCQIHRVDDIQARDEVEGFLQDTFPQQLEESEKQRLGGDVQSP<br>NCPSEDVVITPESFGRDSSLTCLAGNVSACDAPILSSSRSLDCRESGKNGPH<br>VYQDLLLSLGTTNSTLPPPFSLQSGILTLNPVAQGQPILTSLGSNQEEAYVT<br>MSSFYQNQ |

**Table S2. Kinetic parameters for the minibinders characterized using surface plasmon resonance (SPR).**

| Minibinder ID | $k_{on}$ ($M^{-1}$ $s^{-1}$) | $k_{off}$ ($s^{-1}$) | $K_D$ (nM) | $R_{max}$ (RU) |
|---|---|---|---|---|
| Mb-1 (parent) | - | - | - | - |
| Mb-1 (evolved) | $6.0 \times 10^5 \pm 4.1 \times 10^3$ | $1.3 \times 10^{-3} \pm 1.1 \times 10^{-4}$ | $2.1 \pm 0.2$ | $210.7 \pm 18.1$ |
| Mb-2 (parent) | - | - | - | - |
| Mb-2 (evolved) | $1.6 \times 10^5 \pm 1 \times 10^4$ | $5.4 \times 10^{-3} \pm 5.4 \times 10^{-4}$ | $34.0 \pm 4.0$ | $103.5 \pm 39.5$ |
| Mb-3 (parent) | $2.8 \times 10^5 \pm 0.1 \times 10^5$ | $3.6 \times 10^{-2} \pm 0.09 \times 10^{-2}$ | $129 \pm 5.7$ | $202.2 \pm 2.6$ |
| Mb-3 (evolved) | $2.2 \times 10^5 \pm 2.2 \times 10^3$ | $8.9 \times 10^{-4} \pm 4.1 \times 10^{-5}$ | $4.0 \pm 0.2$ | $577.4 \pm 87.1$ |
| Mb-4 (parent) | $3.8 \times 10^5 \pm 4.2 \times 10^3$ | $3.0 \times 10^{-3} \pm 1.7 \times 10^{-5}$ | $7.8 \pm 0.1$ | $444.7 \pm 1.2$ |
| Mb-4 (evolved) | $2.6 \times 10^5 \pm 3.7 \times 10^4$ | $1.4 \times 10^{-4} \pm 2.7 \times 10^{-6}$ | $0.56 \pm 0.08$ | $439.7 \pm 56.9$ |

**Table S3. Mutations observed in the affinity matured minibinders.** For the "predicted to contact IL-7Rα" entries, the IL-7Rα residue numbers are with respect to the start of the extracellular domain (bolded in Table S2) with the first residue being glutamate (E).

| Minibinder ID | Mutation | Predicted location | Predicted to contact IL-7Rα |
|---|---|---|---|
| Mb-1 | N7D | Linker region between α-helix and β-strand. | Yes. Less than 4 Å from histidine 191. |
| | F10Y | An α-helix contacting IL-7Rα. Adjacent to N7D and A14D in 3-D space. | Yes. Less than 4 Å from tyrosine 192. |
| | A14V | An α-helix contacting IL-7Rα. Adjacent to F10Y in 3-D space. | No. |
| | P29S | A β-strand distal to the alpha helices contacting IL-7Rα. | No. |
| | K45E | An α-helix contacting IL-7Rα, with side chain facing away from IL-7Rα. | No. |
| | K52E | An α-helix contacting IL-7Rα, with side chain facing away from IL-7Rα. | No. |
| Mb-2 | L29S | An α-helix not predicted to contact IL-7Rα. Adjacent to V32I in 3-D space. | No. |
| | V32I | An α-helix not predicted to contact IL-7Rα. Adjacent to L29S in 3-D space. | No. |
| | E46K | An α-helix contacting IL-7Rα, with side chain not facing toward IL-7Rα. | No. |
| | E56S | An α-helix contacting IL-7Rα, with side chain facing away from IL-7Rα. Adjacent to A57T in linear sequence and 3-D space. | No. |
| | A57T | An α-helix contacting IL-7Rα. Adjacent to E56S in linear sequence and 3-D space. | No. |
| Mb-3 | M10I | An α-helix contacting IL-7Rα. | Yes. Less than 4 Å from isoleucine 82. |
| | D26G | An α-helix contacting IL-7Rα. | Yes. Less than 4 Å from serine 31 and lysine 138. |
| | K41E | Linker region connecting two α-helices, with side chain facing away from IL-7Rα. | No. |
| Mb-4 | A13T | Linker region connecting an α-helix which contacts IL-7Rα, and a beta-strand. | No. |
| | A24V | An α-helix contacting IL-7Rα. | Proximal to phenylalanine 79 |
| | F25P | An α-helix contacting IL-7Rα. | Yes. Less than 4 Å from leucine 80. |

| | | | |
|---|---|---|---|
| | G40D | Flexible linker connecting two β-strands that do not contact IL-7Rα. | No. |

**Table S4. Yeast strains used in this study.**

| Yeast strain name | Description | Growth medium | Genotype |
|---|---|---|---|
| yAP174 | β-estradiol inducible yeast surface display strain. Used for minibinder expression from CEN/ARS nuclear plasmids. | SC-HU +Hygromycin B (optional due to genomic integration) | MATa AGA1::pMAA29(pER-AGA1-HygR) ura3-52:: pRW122(synTF-URA3) trp1Δ leu2delta1 his3delta200 pep4::HIS3 prb1delta1.6R can1 GAL |
| yAP196 | β-estradiol inducible AHEAD strain containing a p1 landing pad and BadBoy3 error prone DNA polymerase encoded on a CEN/ARS nuclear plasmid. Used to affinity mature and diversify the minibinders. | SC-HLUMC +Hygromycin B (optional due to genomic integration) | MATa AGA1::pMAA29(pER-AGA1-HygR) ura3-52::pRW122(synTF-URA3) trp1Δ leu2delta1 his3delta200 pep4::HIS3 prb1delta1.6R can1 GAL met15Δ; p1-MET15 landing pad; pAW729 |
| EBY100 | Galactose inducible yeast surface display strain. | SC-U | MATa AGA1::GAL1-AGA1::URA3 ura3-52 trp1 leu2-delta200 his3-delta200 pep4::HIS3 prbd1.6R can1 GAL |

**Table S5. Plasmids used in this study.**

| **Plasmid name** | **Description** | **DNA sequence** |
|---|---|---|
| pMAA29 | Used to replace the native *AGA1* locus with an *AGA1* gene controlled a β-estradiol inducible promoter (pER). Contains a Hygromycin resistance cassette. Digest with EcoRI to linearize in preparation for genomic integration. | AAAAGGATCTCAAGAAGATCCTTTGATCTTTTCTACGGGGTCTGACGCTCAGTGGAACGAAAACTCACGTTAAGGGATTTTGGTCA<br>TGAGATTATCAAAAAGGATCTTCACCTAGATCCTTTTAAATTAAAAATGAAGTTTTAAATCAATCTAAAGTATATATGAGTAAACTT<br>GGTCTGACAGTTACCAATGCTTAATCAGTGAGGCACCTATCTCAGCGATCTGTCTATTTCGTTCATCCATAGTTGCCTGACTCCCCG<br>TCGTGTAGATAACTACGATACGGGAGGGCTTACCATCTGGCCCCAGTGCTGCAATGATACCGCGAGACCCACGCTCACCGGCTCCA<br>GATTTATCAGCAATAAACCAGCCAGCCGGAAGGGCCGAGCGCAGAAGTGGTCCTGCAACTTTATCCGCCTCCATCCAGTCTATTAA<br>TTGTTGCCGGGAAGCTAGAGTAAGTAGTTCGCCAGTTAATAGTTTGCGCAACGTTGTTGCCATTGCTACAGGCATCGTGGTGTCAC<br>GCTCGTCGTTTGGTATGGCTTCATTCAGCTCCGGTTCCCAACGATCAAGGCGAGTTACATGATCCCCCATGTTGTGCAAAAAAGCG<br>GTTAGCTCCTTCGGTCCTCCGATCGTTGTCAGAAGTAAGTTGGCCGCAGTGTTATCACTCATGGTTATGGCAGCACTGCATAATTCT<br>CTTACTGTCATGCCATCCGTAAGATGCTTTTCTGTGACTGGTGAGTACTCAACCAAGTCATTCTGAGAATAGTGTATGCGGCGACCG<br>AGTTGCTCTTGCCCGGCGTCAATACGGGATAATACCGCGCCACATAGCAGAACTTTAAAAGTGCTCATCATTGGAAAACGTTCTTC<br>GGGGCGAAAACTCTCAAGGATCTTACCGCTGTTGAGATCCAGTTCGATGTAACCCACTCGTGCACCCAACTGATCTTCAGCATCTTT<br>TACTTTCACCAGCGTTTCTGGGTGAGCAAAAACAGGAAGGCAAAATGCCGCAAAAAAGGGAATAAGGGCGACACGGAAATGTTG<br>AATACTCATACTCTTCCTTTTTCAATATTATTGAAGCATTTATCAGGGTTATTGTCTCATGAGCGGATACATATTTGAATGTATTTAG<br>AAAAATAAACAAATAGGGGTTCCGCGCACATTTCCCCGAAAAGTGCCACCTGACGTCGACGGATCGGGAGATCTCCCGATCCCCT<br>ATGGTGCACTCTCAGTACAATCTGCTCTGATGCCGCATAGTTAAGCCAGTATCTGCTCCCTGCTTGTGTGTTGGAGGTCGCTGAGTA<br>GTGCGCGAGCAAAATTTAAGCTACAACAAGGCAAGGCTTGACCGACAATTGCATGAAGAATCTGCTTAGGGTTAGGCGTTTTGCGC<br>TGCTTCGCGATGTACGGGCCAGATATACGCGTTGgtgagtgaattcAGGTCCAAATGCTGTGTTGTACTACTTAGCTGAAAATGAGTCGAA<br>ACTCTTTGCATTTATCTATAAATTGTTTGGCTCTGTTCCTGGATGGGACAAGAAATTTACTACTGAGCAGCTTGAGGCTTTCAACCA<br>TCAATTTGAATCATCTAACTTTACTGCACGTGAATTGGATCTTGAGTTGCAAAAGGATGTTGCCAGAGTGATTTTAACTCAAGTCGG<br>TTCAGGCCCACAAGAAACAAACGAATCTTTGATTGACGCAAAGACTGGTCTACCAAAGGAATAAGATCAATTCGATATGTAACATT<br>TTTCTTTTCTTTTCTTTTCCTTTTTTTACAATAGCTAATTTACGTTTCCCTACGGTATTGGTCGGAACGACCAAGCTTCAATTTATAAA<br>TATCTTAATTTTAACAGCAGTTACCACTTGAATGAGAAACTTTTTGATGCCTACTCTTTTGCTTTTTCATATCATGACCTCCCTACAT<br>GCTTCGGAATTTCCTTATagcttgcctcgtccccgccgggtcacccggccagcgacatggaggcccagaataccctccttgacagtcttgacgtgcgcagctcaggggcatgatgtgactgtcgccc<br>gtacatttagcccatacatccccatgtataatcatttgcatccatacattttgatggccgcacggcgcgaagcaaaaattacggctcctcgctccagacctgcgagcagggaaacgctcccctcacagacgcgttgaattg<br>tccccacgccgcgcccctgtagagaaatataaaaggttaggatttgccactgaggttcttctttcatatacttccttttaaaatcttgctaggatacagttctcacatcacatccgaacataaacaaaaatgggtaaaaagcct<br>gaactcaccgcgacgtctgtcgagaagtttctgatcgaaaagttcgacagcgtgtccgacctgatgcagctctcggagggcgaagaatctcgtgctttcagcttcgatgtaggagggcgtggatatgtcctgcgggtaa<br>atagctgcgccgatggtttctacaaagatcgttatgtttatcggcactttgcatcggccgcgctcccgattccggaagtgcttgacattggggaatttagcgagagcctgacctattgcatctcccgccgtgcacagggtgt<br>cacgttgcaagacctgcctgaaaccgaactgcccgctgttctgcaaccggtcgcggaggccatggatgcgatcgctgcggccgatcttagccagacgagcgggttcggcccattcggaccgcaaggaatcggtcaa<br>tacactacatggcgtgatttcatatgcgcgattgctgatccccatgtgtatcactggcaaactgtgatggacgacaccgtcagtgcgtccgtcgcgcaggctctcgatgagctgatgctttgggccgaggactgccccga<br>agtccggcacctcgtgcacgcggatttcggctccaacaatgtcctgacggacaatggccgcataacagcggtcattgactggagcgaggcgatgttcggggattcccaatacgaggtcgccaacatcttcttctggag<br>gccgtggttggcttgtatggagcagcagacgcgctacttcgagcggaggcatccggagcttgcaggatcgccgcggctccgggcgtatatgctccgcattggtcttgaccaactctatcagagcttggttgacggcaat<br>ttcgatgatgcagcttgggcgcagggtcgatgcgacgcaatcgtccgatccggagccgggactgtcgggcgtacacaaatcgcccgcagaagcgcggccgtctggaccgatggctgtgtagaagtactcgccgat<br>agtggaaaccgacgccccagcactcgtccgagggcaaaggaataaagtaactgacaataaaaagattcttgttttcaagaacttgtcatttgtatagtttttttatattgtagttgttctattttaatcaaatgttagcgtgatttat<br>atttttttcgcctcgacatcatctgcccagatgcgaagttaagtgcgcagaaagtaatatcatgcgtcaatcgtatgtgaatgctggtcgctatactgcagttcgagtttaAACGCCTAGGTTGAGGCCAA<br>CGaTTATGGGAGatcgaatattaataTTATGGGAGatcgaatattaataTTATGGGAGatcgaatattaataTTATGGGAGatcgaatattaataTTATGGGAGatcgaatatta<br>ataTTATGGGAGatcgaatattaataTTATGGGAGaTGGCATGAATGGCCTTAAGcagatccgccaggcgtgtatatatagcgtggatggccaggcaactttagtgctgacacataca<br>ggcatatatatatgtgtgcgacgacacatgatcatatggcatgcatgtgctctgtatgtatataaaactcttgttttcttcttttctctaaatattctttccttatacattaggacctttgcagcataaattactatacttctatagacaca<br>caaacacaaatacacacactaaATGACATTATCTTTCGCTCATTTTACCTACCTGTTCACAATATTGTTGGGATTAACTAATATTGCCTTGGCATC<br>TGATCCAGAAACGATTCTAGTGACGATAACCAAGACAAACGATGCAAATGGGGTTGTTACAACTACAGTTTCACCCGCGCTAGTCT<br>CCACATCCACTATCGTTCAAGCTGGCACTACGACATTGTATACGACTTGGTGTCCATTGACGGTATCCACTTCATCTGCTGCCGAAA<br>TAAGTCCTTCAATATCGTACGCTACTACCCTATCCAGATTTAGTACTTTGACATTATCTACAGAAGTCTGCTCCCATGAGGCATGTC<br>CTTCGTCATCGACGTTGCCAACCACCACCTTATCTGTGACTTCCAAGTTCACTTCATATATTTGCCCTACTTGTCACACAACCGCTAT<br>CAGCTCATTATCCGAAGTAGGAACTACAACCGTGGTATCATCCAGCGCCATTGAACCATCAAGTGCCTCTATAATCTCACCTGTCA<br>CCTCTACACTTTCGAGTACAACATCGTCCAATCCAACTACTACCTCCCTAAGTTCGACATCTACATCTCCAAGCTCTACATCTACAT<br>CTCCAAGCTCTACATCTACCTCATCAAGTTCGACATCTACCTCATCAAGTTCGACATCTACCTCATCAAGTTCGACATCTACATCTC<br>CAAGTTCGACATCCACATCTTCAAGTTTGACATCCACATCTTCAAGTTCTACATCTACATCCCAAAGTTCTACATCTACCTCATCAA<br>GTTCGACATCTACATCTCCAAGCTCTACATCTACCTCATCAAGTTCAACATCTACATCTCCAAGTTCTAAATCTACTTCTGCAAGCTC<br>CACTTCCACTTCTTCATATTCAACATCTACATCCCCAAGTTTGACTTCTTCATCTCCAACTTTGGCTTCCACTTCTCCAAGTTCAACA<br>TCTATTAGCTCTACTTTTACTGATTCAACTTCATCCCTTGGCTCCTCTATAGCATCTTCATCAACGTCTGTGTCATTATACAGCCCAT<br>CCACACCTGTTTACTCCGTCCCTTCGACTTCGTCAAATGTTGCAACTCCTTCTATGACTTCTTCAACTGTTGAAACAACTGTTAGTTC<br>ACAAAGTTCGTCTGAATATATCACCAAATCCTCAATTTCTACTACTATCCCATCATTTTCCATGTCTACATATTTCACCACTGTTAGT<br>GGAGTCACTACAATGTATACGACATGGTGTCCTTATAGCTCTGAATCTGAGACTAGCACATTAACCAGTATGCATGAAACGGTTAC<br>AACAGACGCTACAGTCTGCACTCACGAGTCTTGCATGCCCTCGCAGACAACAAGTTTGATTACATCTTCTATAAAAATGTCCACTA<br>AAAACGTCGCAACTTCTGTAAGCACCTCAACGGTTGAATCCTCATATGCATGCTCCACATGTGCTGAAACGTCACACTCGTATTCTT<br>CCGTGCAAACAGCTTCATCAAGTTCTGTAACACAGCAGACCACATCCACAAAGAGTTGGGTAAGTTCAATGACAACTTCGGATGAA<br>GATTTCAATAAGCACGCTACCGGTAAGTATCATGTAACATCTTCAGGTACCTCAACCATTTCGACTAGTGTAAGTGAAGCCACGAG<br>TACATCAAGCATTGACTCAGAATCTCAAGAACAATCATCACACTTATTATCGACATCGGTCCTTTCATCCTCCTCCTTGTCTGCTAC<br>ATTATCCTCTGACAGTACTATTTTGCTATTCAGTTCTGTATCATCACTAAGTGTCGAACAGTCACCAGTTACCACACTTCAAATTTCT<br>TCAACATCAGAGATTTTACAACCCACTTCTTCCACAGCTATTGCTACAATATCTGCCTCTACATCATCACTTTCCGCAACATCTATCT<br>CTACACCATCTACCTCTGTGGAATCGACTATTGAATCTTCATCATTGACTCCGACGGTATCTTCTATTTTCCTCTCATCATCATCTGC<br>TCCCTCTTCTCTACAAACATCTGTTACCACTACAGAAGTTTCCACTACTTCAATCTCCATACAATACCAAACTTCATCAATGGTAAC<br>AATTAGCCAATATATGGGCAGTGGATCGCAAACGCGTTTGCCATTAGGAAAGTTGGTCTTCGCCATCATGGCAGTTGCTTGCAATG<br>TAATTTTCAGTTAATATCAGAACAATTTAATTCATTTATATAAACATATACAATCAATCGAATAAAAATACAACAATGTAATGAAA<br>ACGCTCTCTATTGATATGTATTTGCTGACGCTTATAAATTTATTAAAGTTTTATTTTAAACCGTTCTAGTGACTTCACGCGGGGAGAT<br>ACATCCAATAGTAAATTACAAAATAAGGAAAGTATAAAACAgaattcCCGTCGACCTCTAGCTAGAGCTTGGCGTAATCATGGTCATA<br>GCTGTTTCCTGTGTGAAATTGTTATCCGCTCACAATTCCACACAACATACGAGCCGGAAGCATAAAGTGTAAAGCCTGGGGTGCCT<br>AATGAGTGAGCTAACTCACATTAATTGCGTTGCGCTCACTGCCCGCTTTCCAGTCGGGAAACCTGTCGTGCCAGCTGCATTAATGA<br>ATCGGCCAACGCGCGGGGAGAGGCGGTTTGCGTATTGGGCGCTCTTCCGCTTCCTCGCTCACTGACTCGCTGCGCTCGGTCGTTCGG<br>CTGCGGCGAGCGGTATCAGCTCACTCAAAGGCGGTAATACGGTTATCCACAGAATCAGGGGATAACGCAGGAAAGAACATGTGAG<br>CAAAAGGCCAGCAAAAGGCCAGGAACCGTAAAAAGGCCGCGTTGCTGGCGTTTTTCCATAGGCTCCGCCCCCCTGACGAGCATCA<br>CAAAAATCGACGCTCAAGTCAGAGGTGGCGAAACCCGACAGGACTATAAAGATACCAGGCGTTTCCCCCTGGAAGCTCCCTCGTG<br>CGCTCTCCTGTTCCGACCCTGCCGCTTACCGGATACCTGTCCGCCTTTCTCCCTTCGGGAAGCGTGGCGCTTTCTCAATGCTCACGCT<br>GTAGGTATCTCAGTTCGGTGTAGGTCGTTCGCTCCAAGCTGGGCTGTGTGCACGAACCCCCCGTTCAGCCCGACCGCTGCGCCTTAT<br>CCGGTAACTATCGTCTTGAGTCCAACCCGGTAAGACACGACTTATCGCCACTGGCAGCAGCCACTGGTAACAGGATTAGCAGAGCG<br>AGGTATGTAGGCGGTGCTACAGAGTTCTTGAAGTGGTGGCCTAACTACGGCTACACTAGAAGGACAGTATTTGGTATCTGCGCTCT<br>GCTGAAGCCAGTTACCTTCGGAAAAAGAGTTGGTAGCTCTTGATCCGGCAAACAAACCACCGCTGGTAGCGGTGGTTTTTTTGTTT<br>GCAAGCAGCAGATTACGCGCAGAAA |
| pRW122 | Used to genomically integrate the synthetic transcription factor (synTF; pTEF1-ZF97-4-ER-VP16) which induces expression from pER in the presence | Ttgagatcctttttttctgcgcgtaatctgctgcttgcaaacaaaaaaaccaccgctaccagcggtggtttgtttgccggatcaagagctaccaactctttttccgaaggtaactggcttcagcagagcgcagataccaaat<br>actgttcttctagtgtagccgtagttaggccaccacttcaagaactctgtagcaccgcctacatacctcgctctgctaatcctgttaccagtggctgctgccagtggcgataagtcgtgtcttaccgggttggactcaagacg<br>atagttaccggataaggcgcagcggtcgggctgaacggggggttcgtgcacacagcccagcttggagcgaacgacctacaccgaactgagatacctacagcgtgagctatgagaaagcgccacgcttcccgaag<br>ggagaaaggcggacaggtatccggtaagcggcagggtcggaacaggagagcgcacgagggagcttccagggggaaacgcctggtatctttatagtcctgtcgggtttcgccacctctgacttgagcgtcgatttttg<br>tgatgctcgtcaggggggcggagcctatggaaaaacgccagcaacgcggcctttttacggttcctggccttttgctggccttttgctcacatgttctttcctgcgttatcccctgattctgtggataaccgtgcggccgccaa<br>tgctaaattcgagtgaaacacaggaagatcagaaaatcctcatttcatccatattaacaataatttcaaatgtttatttgcattatttgaaactaggcaagacaagcaacgaaacgtttttgaaaattttgagtattttcaataaattt<br>gtagaggactcagatattgaaaaaaagctacagcaattaatacttgataagaagagtattgagaagggcaacggttcatcatctcatggatctgcacatgaacaaacaccagagtcaaacgacgttgaaattgaggctac<br>tgcgccaattgatgacaatacagacgatgataacaaaccgaagttatctgatgtagaaaaggattaaagatgctaagagatagtgatgatatttcataaataatgtaattctatatatgttaattacctttttttgcgaggcatattt<br>atggtgaaggataagttttgaccatcaaagaaggttaatgtggctgtggtttcagggtccataaagcccacatggataacattacccctgaattcgcatctagactgatgagacgtggtagagccacaaacagccggtac<br>aagcaacgatctccaggaccatctgaatcatgcgcggatgacacgaactcacgacggcgatcacagacattaacccacagtacagacactgcgacaacgtggcaattcgtcgcaatacaacgccttgccaacaggg<br>agttcttcagagacatggaggctcaaaacgaaattattgacagcctagacatcaatagtcatacaacagaaagcgaccacccaactttggctgataatagcgtataaacaatgcatactttgtacgttcaaaatacaatgca<br>gtagatatatttatgcatattacatataatacatatcacataggaagcaacaggcgcgttggacttttaattttcgaggaccgcgaatccttacatcacacccaatcccccacaagtgatcccccacacaccatagcttcaaa<br>atgtttctactcctttttttactcttccagattttctcggactccgcgcatcgccgtaccacttcaaaacacccaagcacagcatactaaatttcccctctttcttcctctagggtgtcgttaattacccgtactaaaggtttggaaaa<br>gaaaaaagacaccgcctcgtttctttttcttcgtcgaaaaaggcaataaaaatttttatcacgtttctttttcttgaaaatttttttttttgattttttttctctttcgatgacctcccattgatatttaagttaataaacggtcatcaatttctca<br>agtttcagtttcatttttcttgttctattacaactttttttacttcttgctcattagaaagaaagcatagcaatctaatctaagttttaattacaaaagatctatgCTCGAGAAAAAAatggagcgtcccttccaatgtagaa<br>tttgcatgagaaacttttcgagacaatctaatttgtcaagacatacccgtactcataccggtgaaaaaccgtttcagtgtcggatatgtatgcgaaatttctcaagaaatgaacatttggttttgcatctacgtacgcacaccgg<br>cgagaagccattccaatgcagaatatgtatgagaaactttagtcaaaaaactggtttgagagttcacctaaaaacacatctgagaggtgatggtgctggattgattgatATCGATtctgctggtgacatgagagctgc<br>aaacctttggccaagcccgttaatgattaaacgatctaagaaaaacagtttagctttgtccctgacggctgatcaaatggtaagtgccttgttggatgctgaacctccaatactctattcagaatatgatcctacaagacctttc<br>agtgaagcttcgatgatgggcttacttactaacttagcagacagggagttagttcacatgatcaactgggcaaaaagggtaccaggttttgttgatttgaccttacatgatcaagtccaccttctagaatgtgcctggctaga<br>aatcctaatgattggacttgtctggcgctcaatggaacacccagtgaagctactttttgctcctaatttgttattggacaggaatcaaggaaaatgtgtagagggcatggttgagattttcgacatgttgttagctacatcatcta<br>gattccgcatgatgaatctgcaaggtgaagagtttgtgtgcctcaaatctattattttgcttaattctggtgtgtacacatttttgtccagcactctgaagtctctggaagaaaaagatcatattcatagagtcctggacaagatc |

| | | |
|---|---|---|
| | of β-estradiol into the URA3 locus. Contains a functional URA3 gene for prototrophic selection. Digest with NotI to linearize in preparation for genomic integration. | [illegible] |
| pAW729 | CEN/ARS plasmid containing the BadBoy3 error prone polymerase controlled by a REV1 promoter. Contains a functional LEU2 gene for prototrophic selection. | [illegible] |
| pMAA27 | Donor plasmid with GFP dropout for installing minibinders fused to AGA2 onto p1. Once the GFP dropout has been replaced with a minibinder (e.g., with Gibson Assembly), the | [illegible] |

| | | |
|---|---|---|
| | plasmid can be linearized with ScaI in preparation for integration onto the p1 landing pad in strain yAP196. Contains a functional TRP1 gene for prototrophic selection. | ttgctggccttttgctcacatgttctttcctgcgttatcccctgattctgtggataaccgtattaccgcctttgagtgagctgataccgctcgccgcagccgaacgaccgagcgcagcgagtcagtgagcgaggaagcgga<br>agagcgcccaatacgcaaaccgcctctccccgcgcgttggccgattcattaatgcagctggcacgacaggtttcccgactggaaagcgggcagtgagcgcaacgcaatagtactataatatatgaattacattattaatt<br>taaaaacTcgtgatgacctatacataggaagatctatagaaacaaaaagattaataactttcaaatatcagaaaaatatagaaacatgtgataagctcatagacatAtaaaaaTATGAGATTCCCATCTA<br>TCTTCACCGCTGTTTTGTTCGCTGCTTCTTCTGCTTTGGCTGCTCCAGCTAACACCACCACCGAAGACGAAACCGCTCAAATCCCAG<br>CTGAAGCTGTTATCGACTACTCTGACTTGGAAGGTGACTTCGACGCTGCTGCTTTGCCATTGTCTAACTCTACCAACAACGGTTTGT<br>CTTCTACCAACACCACCATCGCTTCTATCGCTGCTAAGGAAGAAGGTGTTCAATTGGACAAGAGAGAAGCTGACGCAgagaccgaaagtg<br>aaacgtgatttcatgcgtcattttgaacattttgtaaatcttatttaataatgtgtgcggcaattcacatttaatttatgaatgttttcttaacatcgcggcaactcaagaaacggcaggttcggatcttagctactagagaaagag<br>gagaaatactagatgcgtaaaggcgaagagctgttcactggtgtcgtccctattctggtggaactggatggtgatgtcaacggtcataagttttccgtgcgtggcgagggtgaaggtgacgcaactaatggtaaactgac<br>gctgaagttcatctgtactactggtaaactgccggttccttggccgactctggtaacgacgctgacttatggtgttcagtgctttgctcgttatccggaccatatgaagcagcatgacttcttcaagtccgccatgccggaag<br>gctatgtgcaggaacgcacgatttcctttaaggatgacggcacgtacaaaacgcgtgcggaagtgaaatttgaaggcgataccctggtaaaccgcattgagctgaaaggcattgactttaaagaggacggcaatatcct<br>gggccataagctggaatacaattttaacagccacaatgtttacatcaccgccgataaacaaaaaaatggcattaaagcgaattttaaaattcgccacaacgtggaggatggcagcgtgcagctggctgatcactaccag<br>caaaacactccaatcggtgatggtcctgttctgctgccagacaatcactatctgagcacgcaaagcgttctgtctaaagatccgaacgagaaacgcgatcatatggttctgctggagttcgtaaccgcagcgggcatcac<br>gcatggtatggatgaactgtacaaatgaccaggcatcaaataaaacgaaaggctcagtcgaaagactgggcctttcgttttatctgttgtttgtcggtgaacgctctctactagagtcacactggctcaccttcgggtgggc<br>ctttctgcgtttataggtctcGGTTCCaaggacaatagctcgacgattgaaggtagatacccatacgacgttccagactacgctctgcaggctagtggtggtggtggttctggtggtggtggttctggtggtggtggtt<br>ctgctagcgacgtcCAGGAACTGACAACTATATGCGAGCAAATCCCCTCACCAACTTTAGAATCGACGCCGTACTCTTTGTCAACGACTACT<br>ATTTTGGCCAACGGGAAGGCAATGCAAGGAGTTTTTGAATATTACAAATCAGTAACGTTTGTCAGTAATTGCGGTTCTCACCCCTC<br>AACAACTAGCAAAGGCAGCCCCATAAACACACAGTATGTTTTTTGAatcctaataatGGCCGCAAAAAAAAAAAAAAAAAAAAAAAAAAA<br>AAAAAAAAAAAAAAAAAAAAAAAAAAAAAACCTGTCACCGGATGTGTTTTCCGGTCTGATGAGTC<br>CGTGAGGACGAAACaggactag |
| pMAA28 | CEN/ARS plasmid with GFP dropout for yeast surface display of minibinders. GFP dropout should be replaced with minibinders (e.g., using Golden Gate Assembly) in preparation for surface display on yAP174. Contains a functional LEU2 gene for prototrophic selection. | Tcgaggagaacttctagtatatctacatacctaatattattgccttattaaaaatggaatcccaacaattacatcaaaatccacattctcttcaaaatcaattgtcctgtacttccttgttcatgtgtgttcaaaaacgttatatttata<br>ggataattatactctatttctcaacaagtaattggttgtttggccgagcggtctaaggcgcctgattcaagaaatatcttgaccgcagttaactgtgggaatactcaggtatcgtaagatgcaagagttcgaatctcttagcaa<br>ccattatttttttcctcaacataacgagaacacacaggggcgctatcgcacagaatcaaattcgatgactggaaattttttgttaatttcagaggtcgcctgacgcatataccttttttcaactgaaaaattgggagaaaaagga<br>aaggtgagagcgccggaaccggcttttcatatagaatagagaagcgttcatgactaaatgcttgcatcacaatacttgaagttgacaatattatttaaggacctattgtttttccaataggtggttagcaatcgtcttactttcta<br>acttttcttaccttttacatttcagcaatatatatatatatatttcaaggatataccattctaatgtctgcccctaagaagatcgtcgttttgccaggtgaccacgttggtcaagaaatcacagccgaagccattaaggttcttaaag<br>ctatttctgatgttcgttccaatgtcaagttcgatttcgaaaatcatttaattggtggtgctgctatcgatgctacaggtgttccacttccagatgaggcgctggaagcctccaagaaggctgatgccgttttgttaggtgctgtg<br>ggtggtcctaaatggggaaccggtagtgttagacctgaacaaggtttactaaaaatccgtaaagaacttcaattgtacgccaacttaagaccatgtaactttgcatccgactctcttttagacttatctccaatcaagccacaa<br>tttgctaaaggtactgacttcgttgttgtcagagaattagtgggaggtatttactttggtaagagaaaggaggacgatggtgatggtgtcgcttgggatagtgaacaatacaccgttccagaagtgcaaagaatcacaaga<br>atggccgctttcatggccctacaacatgagccaccattgcctatttggtccttggataaagctaatgttttggcctcttcaagattatggagaaaaactgtggaggaaaccatcaagaacgaatttcctacattgaaggttcaa<br>catcaattgattgattctgccgccatgatcctagttaagaacccaacccacctaaatggtattataatcaccagcaacatgtttggtgatatcatctccgatgaagcctccgttatcccaggttccttgggtttgttgccatctgc<br>gtccttggcctctttgccagacaagaacaccgcatttggtttgtacgaaccatgccacggttctgctccagatttgccaaagaataaggtcaaccctatcgccactatcttgtctgctgcaatgatgttgaaattgtcattgaa<br>cttgcctgaagaaggtaaggccattgaagatgcagttaaaaaggttttggatgcaggtatcagaactggtgatttaggtggttccaacagtaccaccgaagtcggtgatgctgtcgccgaagaagttaagaaaatccttg<br>cttaacctggaggacccttctctttagactattctactcttatgcacgtaaaaaattctaggaaatatgtattaactaggagtaaaataaccggctagtggcattcatatagccgtctgtttacatctacatcacacatttcgagtg<br>tatatctcgcaacgttggcgttaaataggcagatcacgtgctataaaaataattataatttaaatttttaatataaatatataaattaaaaatagaaagtaaaaaaagaaattaaagaaaaaatagtttttgttttccgaagatgta<br>aaagactctagggggatcgccaacaaatactaccttttatcttgctcttcctgctctcaggtattaatgccgaattgtttcatcttgtctgtgtagaaaaccacacacgaaaatcctgtgattttacattttacttatcgttaatcgaa<br>tgtatatctatttaatctgcttttcttgtctaataaatatatatgtaaagtacgctttttgttgaaattttttaaacctttgtttattttttttttcttcattccgtaactcttctaccttctttatttactttctaaaatccaaatacaaaacataaaaa<br>taaataaacacagagtaaattcccaaattattccatcattaaaagatacgaggcgcgtgtaagttacaggcaagcgatccgtcccgagcggccgcgattatcaaaaaggatcttcacctagatccttttaaattaaaaatga<br>agttttaaatcaatctaaagtatatatgagtaaacttggtctgacagttaccaatgcttaatcagtgaggcacctatctcagcgatctgtctatttcgttcatccatagttgcctgactccccgtcgtgtagataactacgatacgg<br>gagggcttaccatctggccccagtgctgcaatgataccgcgggacccacgctcaccggctccagatttatcagcaataaaccagccagccggaagggccgagcgcagaagtggtcctgcaactttatccgcctccat<br>ccagtctattaattgttgccgggaagctagagtaagtagttcgccagttaatagtttgcgcaacgttgttgccattgctacaggcatcgtggtgtcacgctcgtcgtttggtatggcttcattcagctccggttcccaacgatca<br>aggcgagttacatgatcccccatgttgtgcaaaaaagcggttagctccttcggtcctccgatcgttgtcagaagtaagttggccgcagtgttatcactcatggttatggcagcactgcataattctcttactgtcatgccatcc<br>gtaagatgcttttctgtgactggtgagtactcaaccaagtcattctgagaatagtgtatgcggcgaccgagttgctcttgcccggcgtcaatacgggataataccgcgccacatagcagaactttaaaagtgctcatcattg<br>gaaaacgttcttcggggcgaaaactctcaaggatcttaccgctgttgagatccagttcgatgtaacccactcgtgcacccaactgatcttcagcatcttttactttcaccagcgtttctgggtgagcaaaaacaggaaggca<br>aaatgccgcaaaaaagggaataagggcgacacggaaatgttgaatactcatactcttcctttttcaatattattgaagcatttatcagggttattgtctcatgagcggatacatatttgaatgtatttagaaaaataaacaaata<br>ggggttccgcgcacatttccccgaaaagtgccacctgtcatgaccaaaatcccttaacgtgagttttcgttccactgagcgtcagaccccgtagaaaagatcaaaggatcttcttgagatcctttttttctgcgcgtaatctgc<br>tgcttgcaaacaaaaaaaccaccgctaccagcggtggtttgtttgccggatcaagagctaccaactctttttccgaaggtaactggcttcagcagagcgcagataccaaatactgttcttctagtgtagccgtagttaggcc<br>accacttcaagaactctgtagcaccgcctacatacctcgctctgctaatcctgttaccagtggctgctgccagtggcgataagtcgtgtcttaccgggttggactcaagacgatagttaccggataaggcgcagcggtcg<br>ggctgaacggggggttcgtgcacacagcccagcttggagcgaacgacctacaccgaactgagatacctacagcgtgagctatgagaaagcgccacgcttcccgaagggagaaaggcggacaggtatccggtaag<br>cggcagggtcggaacaggagagcgcacgagggagcttccagggggaaacgcctggtatctttatagtcctgtcgggtttcgccacctctgacttgagcgtcgatttttgtgatgctcgtcaggggggcggagcctatg<br>gaaaaacgccagcaacgcggcctttttacggttcctggccttttgctggccttttgctcacatgttctttcctgcgttatcccctgattctgtggataaccgtgcggccgcccctgaattcgcatctagactgatgagacgtgg<br>tagagccacaaacagccggtacaagcaacgatctccaggaccatctgaatcatgcgcggatgacacgaactcacgacggcgatcacagacattaacccacagtacagacactgcgacaacgtggcaattcgtcgca<br>atacaacgCCTAGGTTGAGGCCAACGaTTATGGGAGatcgaatattaataTTATGGGAGatcgaatattaataTTATGGGAGatcgaatattaataTTATGGGAGatcg<br>aatattaataTTATGGGAGatcgaatattaataTTATGGGAGatcgaatattaataTTATGGGAGaTGGCATGAATGGCCTTAAGcagatccgccaggcgtgtatatatagcgt<br>ggatggccaggcaactttagtgctgacacatacaggcatatatatatgtgtgcgacgacacatgatcatatggcatgcatgtgctctgtatgtatataaaactcttgttttcttcttttctctaaatattctttccttatacattagga<br>cctttgcagcataaattactatacttctatagacacacaaacacaaatacacacactaaATGAGATTCCCATCTATCTTCACCGCTGTTTTGTTCGCTGCTTCTTCTGCTTT<br>GGCTGCTCCAGCTAACACCACCACCGAAGACGAAACCGCTCAAATCCCAGCTGAAGCTGTTATCGACTACTCTGACTTGGAAGGTG<br>ACTTCGACGCTGCTGCTTTGCCATTGTCTAACTCTACCAACAACGGTTTGTCTTCTACCAACACCACCATCGCTTCTATCGCTGCTAA<br>GGAAGAAGGTGTTCAATTGGACAAGAGAGAAGCTGACGCAgagaccgaaagtgaaacgtgatttcatgcgtcattttgaacattttgtaaatcttatttaataatgtgtgcggcaatt<br>cacatttaatttatgaatgttttcttaacatcgcggcaactcaagaaacggcaggttcggatcttagctactagagaaagaggagaaatactagatgcgtaaaggcgaagagctgttcactggtgtcgtccctattctggtg<br>gaactggatggtgatgtcaacggtcataagttttccgtgcgtggcgagggtgaaggtgacgcaactaatggtaaactgacgctgaagttcatctgtactactggtaaactgccggttccttggccgactctggtaacgac<br>gctgacttatggtgttcagtgctttgctcgttatccggaccatatgaagcagcatgacttcttcaagtccgccatgccggaaggctatgtgcaggaacgcacgatttcctttaaggatgacggcacgtacaaaacgcgtgc<br>ggaagtgaaatttgaaggcgataccctggtaaaccgcattgagctgaaaggcattgactttaaagaggacggcaatatcctgggccataagctggaatacaattttaacagccacaatgtttacatcaccgccgataaac<br>aaaaaaatggcattaaagcgaattttaaaattcgccacaacgtggaggatggcagcgtgcagctggctgatcactaccagcaaaacactccaatcggtgatggtcctgttctgctgccagacaatcactatctgagcacg<br>caaagcgttctgtctaaagatccgaacgagaaacgcgatcatatggttctgctggagttcgtaaccgcagcgggcatcacgcatggtatggatgaactgtacaaatgaccaggcatcaaataaaacgaaaggctcagt<br>cgaaagactgggcctttcgttttatctgttgtttgtcggtgaacgctctctactagagtcacactggctcaccttcgggtgggcctttctgcgtttataggtctcGGTTCCaaggacaatagctcgacgattgaaggtaga<br>tacccatacgacgttccagactacgctctgcaggctagtggtggtggtggttctggtggtggtggttctggtggtggtggttctgctagcgacgtcCAGGAACTGACAACTATATGCGAGCAA<br>ATCCCCTCACCAACTTTAGAATCGACGCCGTACTCTTTGTCAACGACTACTATTTTGGCCAACGGGAAGGCAATGCAAGGAGTTTTT<br>GAATATTACAAATCAGTAACGTTTGTCAGTAATTGCGGTTCTCACCCCTCAACAACTAGCAAAGGCAGCCCCATAAACACACAGTA<br>TGTTTTTTGAtagctcgagatctgataacaacagtgtagatgtaacaaaatcgactttgttcccactgtacttttagctcgtacaaaatacaatatacttttcatttctccgtaaacaacatgttttcccatgtaatatcctt<br>ttctatttttcgttccgttaccaactttacacatactttatatagctattcacttctatacactaaaaaactaagacaattttaattttgctgcctgccatatttcaatttgttataaattcctataatttatcctattagtagctaaaaaaaga<br>tgaatgtgaatcgaatcctaagagaatt |

**Table S6: Oligonucleotides used in this study.** All oligonucleotides were purified by Integrated DNA Technologies (IDT) with standard desalting.

| Oligo ID | Description | |
|---|---|---|
| MAA_o169 | Mb-1 segment: 1<br>Anneals with: MAA_o170<br>Mutations contained: parent sequence | ACGCATGTTGTCTAACTTTTGAAAA CCCTGAGTTCGCCGAACAAGCTGTG CTGATT |
| MAA_o170 | Mb-1 segment: 1<br>Anneals with: MAA_o169<br>Mutations contained: parent sequence | CGGCAATCAGCACAGCTTGTTCGGC GAACTCAGGGTTTTCAAAAGTTAGA CAACAT |
| MAA_o171 | Mb-1 segment: 1<br>Anneals with: MAA_o172<br>Mutations contained: N7D | ACGCATGTTGTCTAACTTTTGAAGA CCCTGAGTTCGCCGAACAAGCTGTG CTGATT |
| MAA_o172 | Mb-1 segment: 1<br>Anneals with: MAA_o171<br>Mutations contained: N7D | CGGCAATCAGCACAGCTTGTTCGGC GAACTCAGGGTCTTCAAAAGTTAGA CAACAT |
| MAA_o173 | Mb-1 segment: 1<br>Anneals with: MAA_o174<br>Mutations contained: F10Y | ACGCATGTTGTCTAACTTTTGAAAA CCCTGAGTACGCCGAACAAGCTGTG CTGATT |
| MAA_o174 | Mb-1 segment: 1<br>Anneals with: MAA_o173<br>Mutations contained: F10Y | CGGCAATCAGCACAGCTTGTTCGGC GTACTCAGGGTTTTCAAAAGTTAGA CAACAT |
| MAA_o175 | Mb-1 segment: 1<br>Anneals with: MAA_o176<br>Mutations contained: A14V | ACGCATGTTGTCTAACTTTTGAAAA CCCTGAGTTCGCCGAACAAGTTGTG CTGATT |
| MAA_o176 | Mb-1 segment: 1<br>Anneals with: MAA_o175<br>Mutations contained: A14V | CGGCAATCAGCACAACTTGTTCGGC GAACTCAGGGTTTTCAAAAGTTAGA CAACAT |
| MAA_o177 | Mb-1 segment: 1<br>Anneals with: MAA_o178<br>Mutations contained: N7D, F10Y | ACGCATGTTGTCTAACTTTTGAAGA CCCTGAGTACGCCGAACAAGCTGTG CTGATT |
| MAA_o178 | Mb-1 segment: 1<br>Anneals with: MAA_o177<br>Mutations contained: N7D, F10Y | CGGCAATCAGCACAGCTTGTTCGGC GTACTCAGGGTCTTCAAAAGTTAGA CAACAT |
| MAA_o179 | Mb-1 segment: 1<br>Anneals with: MAA_o180<br>Mutations contained: N7D, A14V | ACGCATGTTGTCTAACTTTTGAAGA CCCTGAGTTCGCCGAACAAGTTGTG CTGATT |
| MAA_o180 | Mb-1 segment: 1<br>Anneals with: MAA_o179<br>Mutations contained: N7D, A14V | CGGCAATCAGCACAACTTGTTCGGC GAACTCAGGGTCTTCAAAAGTTAGA CAACAT |
| MAA_o181 | Mb-1 segment: 1<br>Anneals with: MAA_o182<br>Mutations contained: F10Y, A14V | ACGCATGTTGTCTAACTTTTGAAAA CCCTGAGTACGCCGAACAAGTTGTG CTGATT |
| MAA_o182 | Mb-1 segment: 1<br>Anneals with: MAA_o181<br>Mutations contained: F10Y, A14V | CGGCAATCAGCACAACTTGTTCGGC GTACTCAGGGTTTTCAAAAGTTAGA CAACAT |
| MAA_o183 | Mb-1 segment: 1<br>Anneals with: MAA_o184<br>Mutations contained: N7D, F10Y, A14V | ACGCATGTTGTCTAACTTTTGAAGA CCCTGAGTACGCCGAACAAGTTGTG CTGATT |

| | | |
|---|---|---|
| MAA_o184 | Mb-1 segment: 1<br>Anneals with: MAA_o183<br>Mutations contained: N7D, F10Y, A14V | CGGCAATCAGCACAACTTGTTCGGC<br>GTACTCAGGGTCTTCAAAAGTTAGA<br>CAACAT |
| MAA_o185 | Mb-1 segment: 2<br>Anneals with: MAA_o186<br>Mutations contained: parent sequence | GCCGAAGATGCTGGCTTGAAAGTCG<br>AGGTAAAGCCAGGTGAAGACT |
| MAA_o186 | Mb-1 segment: 2<br>Anneals with: MAA_o185<br>Mutations contained: parent sequence | AATGAGTCTTCACCTGGCTTTACCTC<br>GACTTTCAAGCCAGCATCTT |
| MAA_o187 | Mb-1 segment: 2<br>Anneals with: MAA_o188<br>Mutations contained: P29S | GCCGAAGATGCTGGCTTGAAAGTCG<br>AGGTAAAGTCAGGTGAAGACT |
| MAA_o188 | Mb-1 segment: 2<br>Anneals with: MAA_o187<br>Mutations contained: P29S | AATGAGTCTTCACCTGACTTTACCTC<br>GACTTTCAAGCCAGCATCTT |
| MAA_o189 | Mb-1 segment: 3<br>Anneals with: MAA_o190<br>Mutations contained: parent sequence | CATTAACCGTTTGCCTTTCTAATGAA<br>GCAGCATGTAAGTATTTTGC |
| MAA_o190 | Mb-1 segment: 3<br>Anneals with: MAA_o189<br>Mutations contained: parent sequence | TTCAGCAAAATACTTACATGCTGCT<br>TCATTAGAAAGGCAAACGGTT |
| MAA_o193 | Mb-1 segment: 3<br>Anneals with: MAA_o194<br>Mutations contained: K45E | CATTAACCGTTTGCCTTTCTAATGAA<br>GCAGCATGTGAGTATTTTGC |
| MAA_o194 | Mb-1 segment: 3<br>Anneals with: MAA_o193<br>Mutations contained: K45E | TTCAGCAAAATACTCACATGCTGCT<br>TCATTAGAAAGGCAAACGGTT |
| MAA_o197 | Mb-1 segment: 4<br>Anneals with: MAA_o198<br>Mutations contained: parent sequence | TGAAAGAGCGAAAGAGATGGGAGT<br>TCCCGTTACAGTCGAACCAG |
| MAA_o198 | Mb-1 segment: 4<br>Anneals with: MAA_o197<br>Mutations contained: parent sequence | GAACCTGGTTCGACTGTAACGGGAA<br>CTCCCATCTCTTCCGCTCT |
| MAA_o199 | Mb-1 segment: 4<br>Anneals with: MAA_o200<br>Mutations contained: K52E | TGAAAGAGCGGAAGAGATGGGAGT<br>TCCCGTTACAGTCGAACCAG |
| MAA_o200 | Mb-1 segment: 4<br>Anneals with: MAA_o199<br>Mutations contained: K52E | GAACCTGGTTCGACTGTAACGGGAA<br>CTCCCATCTCTTTCGCTCT |
| MAA_o254 | Used to amplify the diversified Mb-4 library in preparation for Gibson Assembly. | GGACAAGAGAGAAGCTGACGCA |
| MAA_o255 | Used to amplify the diversified Mb-4 library in preparation for Gibson Assembly. | cgtcgagctattgtccttGGAACC |
| MAA_o256 | Primer used to amplify minibinder variants in bin 1. Contains R1 Illumina adapter.<br>Barcode: **gtca** | ACACTCTTTCCCTACACGACGCTCTT<br>CCGATCT**gtca**GTTCAATTGGACAAG<br>AGAGAAGC |

| | | |
|---|---|---|
| MAA_o257 | Primer used to amplify minibinder variants in bin 1. Contains R2 Illumina adapter.<br>Barcode: **gtca** | GACTGGAGTTCAGACGTGTGCTCTTCCGATCT**gtca**CGTCGAGCTATTGTCCTTGG |
| MAA_o258 | Primer used to amplify minibinder variants in bin 2. Contains R1 Illumina adapter.<br>Barcode: **tcgt** | ACACTCTTTCCCTACACGACGCTCTTCCGATCT**tcgt**GTTCAATTGGACAAGAGAGAAGC |
| MAA_o259 | Primer used to amplify minibinder variants in bin 2. Contains R2 Illumina adapter.<br>Barcode: **tcgt** | GACTGGAGTTCAGACGTGTGCTCTTCCGATCT**tcgt**CGTCGAGCTATTGTCCTTGG |
| MAA_o260 | Primer used to amplify minibinder variants in bin 3. Contains R1 Illumina adapter.<br>Barcode: **caag** | ACACTCTTTCCCTACACGACGCTCTTCCGATCT**caag**GTTCAATTGGACAAGAGAGAAGC |
| MAA_o261 | Primer used to amplify minibinder variants in bin 3. Contains R2 Illumina adapter.<br>Barcode: **caag** | GACTGGAGTTCAGACGTGTGCTCTTCCGATCT**caag**CGTCGAGCTATTGTCCTTGG |
| MAA_o262 | Primer used to amplify minibinder variants in bin 4. Contains R1 Illumina adapter.<br>Barcode: **agtc** | ACACTCTTTCCCTACACGACGCTCTTCCGATCT**agtc**GTTCAATTGGACAAGAGAGAAGC |
| MAA_o263 | Primer used to amplify minibinder variants in bin 4. Contains R2 Illumina adapter.<br>Barcode: **agtc** | GACTGGAGTTCAGACGTGTGCTCTTCCGATCT**agtc**CGTCGAGCTATTGTCCTTGG |
| K_221 | Primer used to amplify sorted minibinder variants from the neutral drift cycles 8, 15, and 30 library. Contains R1 Illumina adapter. | ACACTCTTTCCCTACACGACGCTCTTCCGATCTagctGTTCAATTGGACAAGAGAGAAGC |
| K_222 | Primer used to amplify sorted minibinder variants from the neutral drift cycles 8, 15, and 30 library. Contains R2 Illumina adapter. | GACTGGAGTTCAGACGTGTGCTCTTCCGATCTagctCGTCGAGCTATTGTCCTTGG |
